# IsoVEM: Isotropic Reconstruction for Volume Electron Microscopy Based on Transformer

**DOI:** 10.1101/2023.11.22.567807

**Authors:** Jia He, Yan Zhang, Wenhao Sun, Ge Yang, Fei Sun

**Author notes:** Correspondence: Yan Zhang,; Ge Yang,; and Fei Sun. These authors contributed equally to this work (Jia He, Yan Zhang, Wenhao Sun).

## Abstract

Volume electron microscopy (vEM) has become a rapidly developing technique for studying the 3D architecture of biological specimens, such as cells, tissues, and organs at nanometer resolution; this technique involves collecting a series of electron micrographs of axial sequential sections and reconstructing the 3D volume, providing useful information on the cellular ultrastructural spectrum. This technique currently suffers from anisotropic resolution between the lateral (x, y) and axial (z) directions and the loss/damage of sections. Here, we develop a new algorithm, IsoVEM, based on a video transformer model to boost the axial resolution and achieve isotropic reconstruction of vEM. By learning high-resolution axial structures and utilizing the 3D continuity of biological structures, IsoVEM can recover axial information and repair random lost/damaged sections based on a self-supervision strategy, achieving a higher resolution than existing methods, which has been validated for both simulated FIB-SEM datasets and experimental ssTEM datasets. In addition to visual validation, the segmentation efficiency and statistical precision of various ultrastructures, e.g., neurons, mitochondria, vesicles, and membrane bilayers, also prove the better performance of IsoVEM. Therefore, using IsoVEM, we achieve isotropic reconstruction via anisotropic axial sampling, which increases the vEM throughput for studying large-scale biological architectures.

## 1. Introduction

In recent years, with the continuous development and optimization of electron microscope imaging technology, big data storage, image processing technology and computer hardware technology, volume electron microscopy (vEM) has developed rapidly. vEM allows direct depiction of the 3D ultrastructures of organisms, cells^1-4^ and tissues^5,6^ and has been widely used in connectomic studies of neurons^7-10^. The physical scale studied by vEM is also increasing, from the nanoscale to tens of microns and even hundreds of microns ^9,11^. Thus, the rapid development of vEM has promoted its wide application in the life sciences ^12,13^, medicine ^14-16^ and clinical diagnosis ^17-20^, and vEM has become one of the most important and state-of-the-art technologies in the above fields. For these reasons, vEM was named one of Nature’s top 7 technologies to watch in 2023.

Even with the most advanced technology, there are still difficulties or flaws that need to be overcome for vEM. The different imaging methods used for vEM include volume transmission electron microscopy (TEM) and volume scanning electron microscopy (SEM)^9^. The volume TEM technique includes serial section electron tomography (ssET) and ssTEM. Volume SEM techniques include serial block surface scanning electron microscopy (SBF-SEM)^11^, serial ultrathin section scanning electron microscopy (ssSEM) and focused ion beam scanning electron microscopy (FIB-SEM)^11^. ATUM-SEM ^21,22^ and AutoCUTs-SEM ^23^, which can automatically tape-collect serial sections, are ssSEM methods.

For vEM, the z-axis direction of the reconstructed volumes of ssTEM and ssSEM ^11^ can contain only the integrated information of the thickness of each section, so the section thickness determines the axial voxel size. Usually, the section thickness is approximately tens of nanometers, which is several times or even tens of times the lateral plane thickness; this results in the anisotropy of the information between the lateral and axial directions. Similarly, the axial resolution of SBF-SEM depends on the thickness of the sample sequentially cut by the diamond knife and the depth that the electron beam penetrates. FEI designed the volume scope with automatic deconvolution and multienergy imaging methods to enhance the axial resolution of SBF-SEM further. However, at present, the isotropic resolution achieved by the volume scope is dozens of nanometers. Therefore, ssTEM, ssSEM and SBF-SEM are anisotropic vEM techniques for which the target is ten nanometers or more common. ssET and FIB-SEM can achieve isotropic vEM reconstruction. However, because each section of ssET can be tilted within only a limited angular range, the reconstructed result is not complete in Fourier space because information is lost in the high-angle range; this phenomenon is known as the missing cone problem. At present, none of the methods can fill missing cones with complete accurate high-frequency information. In contrast, ssET can be applied to only small volumes ranging from hundreds of nanometers to several microns. FIB-SEM can be used to control the cutting thickness more precisely by controlling the ion beam current and accelerating voltage to realize isotropic 3D reconstruction. C. Shan Xu and Harald Hess developed enhanced FIB-SEM, which enables large-scale isotropic reconstruction, with cumulative collective volumes up to 10^6^ microns in several months ^24^. However, the time cost is a major issue that must be considered when performing large-volume 3D reconstruction. In addition, enhanced FIB-SEM is available at only a few institutions. Therefore, FIB-SEM is currently applicable mainly to isotropic reconstructions of small-volume samples. However, further development of these techniques is needed for efficient isotropic reconstructions at a large scale and high resolution.

Our research goal is to routinely obtain isotropic high-resolution information from large-volume samples via the above vEM technique. Due to the limitations of the above techniques, large-scale and high-resolution isotropic reconstructions cannot be widely realized. Previously, interpolation was usually applied to reconstruction results to reduce the anisotropy effect caused by the loss of axial information. However, the errors and artifacts caused by interpolation are obvious, especially in the case of anisotropic reconstruction with high-anisotropy factors, such as 8× and 10×.

In recent years, various types of deep learning algorithms have continuously emerged and promoted the development of artificial intelligence. Subsequently, its application in vEM has gradually increased. Heinrich first introduced the concept of deep learning to vEM in an attempt to solve the anisotropic reconstruction problem ^25^, demonstrating the feasibility of 3D-FSRCNN and 3D-SRUNet for performing isotropic reconstruction via supervised learning, and the results are superior to those of bicubic interpolation. However, this was a supervised 3D reconstruction method that was carried out only when ground-truth data were available. However, isotropic ground-truth volumes cannot be obtained via ssSEM/ssTEM/SBF-SEM in vEM studies, which limits the application of such supervised deep learning methods in vEMs. Self-supervised deep learning methods without the need for ground truth are more suitable for isotropic reconstruction of vEMs, including those of Kausumi Hagita et al.^26^, Xue Yang Fu et al.^27^, and Hyoungjun Park et al.^28^ et al. These methods are all based on generating adversarial networks (GANs), using high-resolution two-dimensional images in the lateral direction as unpaired self-supervised signals and guiding the model to restore axial low-resolution images by domain transferring. Specifically, Kausumi Hagita et al.^26^ first used the SRGAN^29^ to restore simulated anisotropic FIB-SEM volumes, and Hyoungjun Park et al.^28^ applied similar methods to fluorescence 3D imaging. Xue Yang Fu et al.^27^ adopted a better CycleGAN^30^ to deal with the blind degradation of vEM and performed isotropic reconstruction on real ssTEM data with a 10× anisotropy factor. However, these GANs still adopt a 2D network architecture, which has shortcomings in modeling the intersection continuity of 3D volumes. And moreover, due to the high difficulty and instability of training, the final reconstruction results may contain many artifacts, such as texture artifacts or structural deformations. Several studies have also used video models to recover axial information; for example, Zejin Wang et al. ^31^ designed a video frame interpolation model with a self-attention mechanism that completes the intermediate section between two sections but achieves only 2× axial information recovery.

More recently, diffusion model-based vEM isotropic reconstruction methods, such as DiffuseIR^32^, EMDiffuse^33^ and DiffusionEM^34^, have emerged. Overall, the diffusion model has better high-frequency information recovery ability and training stability than does the GAN. Among these methods, DiffuseIR and DiffusionEM are used in combination with the zero-shot restoration paradigm; in addition, the diffusion model is trained using lateral sections, after which the anisotropic axial sections are restored during sampling. In contrast, EMDiffuse uses a diffusion model to implement a toolbox for electron microscopy restoration, which includes superresolution, denoising, and isotropic reconstruction. However, these diffusion model methods still use a 2D model architecture, only partially compensating for the lack of spatial continuity through positional encoding. Moreover, sampling and training of diffusion models are time-consuming, which becomes a bottleneck in practical applications. Therefore, it is necessary to design new deep learning models for isotropic reconstruction.

Other factors affecting the effectiveness of vEM reconstruction include artifacts such as section damage, creases, contamination and other human defects, which are inevitable during sample preparation. This not only affects preprocessing operations, such as stitching and alignment between images, but also reduces the overall spatial resolution of the subsequent reconstruction results. For samples with many defects, samples can be reprepared only to obtain high-quality images, which slows the speed and efficiency of vEM research. In recent years, image inpainting methods ^35-39^ involving deep learning have been developed rapidly in image editing and photo restoration, compensating for the disadvantage that the information repaired by traditional methods does not match the original semantics. Moreover, these technologies have also been gradually developed and applied in medical image repair^40^, such as inpainting for MRI spectroscopy images,. Commonly used inpainting networks include convolutional neural networks based on context encoders^41^, fully convolutional neural networks based on global and local context discriminators^42^, vanilla GAN networks based on generating images with random noise^43^, U-Net networks based on pyramid context encoders^44^, transformer networks^45^ and other related methods^46-49^. To our knowledge, the application of inpainting in vEM is still very rare. By using the continuity prior of the ultrastructure between continuous sections or continuous block face images and combining this information with the long-term attention mechanism of the transformer model, we believe that the transformer network can be extended to the inpainting of vEM images with defects such as missing sections, damage, and wrinkles. This approach will not only repair the above defects in vEM raw images but also save time and economic costs in sample preparation. In this work, we propose the IsoVEM deep learning model, which uses a video transformer to solve the isotropic reconstruction problem of vEM and handles image inpainting for defects in the xy plane simultaneously. IsoVEM restores axial information by learning the high-resolution lateral information of anisotropic volumes in a self-supervised manner. By adopting a video transformer architecture that includes self-attention and a mutual attention mechanism, the spatial continuity of 3D structures can be effectively learned, avoiding the discontinuous layered artifacts in 3D reconstruction volumes that typically appear in 2D models. Moreover, for data with section deficiency or even sections lost during data collection, IsoVEM can be extended to IsoVEM+ by performing section image inpainting and isotropic reconstruction simultaneously. For example, IsoVEM+ can achieve reliable section inpainting based on interframe continuity and a video transformer model. In addition, IsoVEM can achieve arbitrary-scale isotropic reconstruction during the inference stage, which enables IsoVEM to be easily transferred to other vEM data with different anisotropic factors, even for fractional factors. The pretrained IsoVEM has good transfer robustness, indicating its potential as a universal pretrained isotropic reconstruction model for vEMs. On both simulated anisotropic FIB-SEM and real anisotropic ssTEM data, IsoVEM significantly improved the morphological restoration of organelle ultrastructures of different types and scales, such as neurons, mitochondria, vesicles and bilayers. Based on multiple qualitative indicators, including reconstruction accuracy, artifact restriction, training stability and speed, IsoVEM is far superior to interpolation, better than existing self-supervised methods, and close to existing supervised methods. This approach further facilitates downstream analysis related to connectomics or structures, including 3D segmentation and statistical analysis.

## 2. Results

For the isotropic reconstruction problem of vEM, we designed a video transformer-based self-supervised deep learning model named IsoVEM (Section 2.1), which shows superior isotropic performance compared with traditional interpolation algorithms and other self-supervised algorithms both for simulated anisotropic FIB-SEM data (Section 2.2) and real anisotropic ssTEM data (Section 2.3). The Fourier shell correlation (FSC) criterion is used to determine the accurate resolution of the reconstructed volume via the IsoVEM. Experiments on simulated anisotropic data demonstrate that for 40 nm section thicknesses and 5 nm lateral spatial sampling, axial information above 18 nm can be accurately recovered. Segmenting organelles such as neurons and mitochondria and the corresponding statistical analysis further demonstrated the accuracy and reliability of IsoVEM. Moreover, IsoVEM can achieve isotropic reconstruction with any anisotropic factor and has good transfer robustness for various vEM data, such as different tissues and organs collected by FIB-SEM, ssTEM or ssSEM (Section 2.4). IsoVEM can be extended to IsoVEM+ when facing common section deficiency, such as when performing section inpainting for missing sections or contaminated or broken areas while isotropic reconstruction occurs (Section 2.5). To the best of our knowledge, IsoVEM is the first method for realizing isotropic reconstruction and inpainting simultaneously.

### 2.1. Algorithm design and network architecture

Since the isotropic ground truth of the vEM cannot be obtained, we adopt a self-supervised learning strategy for calculating the anisotropic volume. For dataset preparation, we crop the anisotropic volume into subvolumes of a specific size to generate training/validation/test datasets (Fig. 1a). The training set usually accounts for 60%∼80%, the validation set is the remaining area that does not overlap with the training set, and the test set can include any other data that are different from those in the training set. To utilize lateral high-resolution information for self-supervision, we perform anisotropic degradation on subvolumes in the training set along the X/Y axis to construct paired training data. The degradation process is performed by an anisotropic average pooling operator (Fig. 1a), which is suitable for using multiple modalities of vEM (Supplementary Note 1) and was used in previous work^25^. During training, the paired subvolumes are augmented with eight types of orthogonal rotation and regarded as video clips along the z-axis (Fig. 1b). After feeding them into IsoVEM, the subvolumes are superresolved along the simulated degradation axis (X/Y), and the weighted sum of the L1 and SSIM differences is taken as the loss function (Fig. 1b). During inference, the input subvolumes are also augmented with eight types of orthogonal rotations and then regarded as video clips along the X/Y axis (Fig. 1b). After model inference, 8 superresolved rotation augmentation results are average ensembled to generate the final reconstruction. Finally, the reconstructed subvolumes in the test dataset are stitched according to the cropping coordinates to obtain the whole isotropic reconstructed volume. The network architecture of IsoVEM (Fig. 1c) is designed as a multiscale symmetrical stack of transformer blocks with an arbitrary-scale upsampling module. Specifically, given a subvolume or video clip, the process flow is as follows. First, feature extraction is performed by spatial 2D convolution for each frame in the video clip. Then, the feature maps are processed by multiple transformer stages, where the 1∼7^th^ stages perform multiscale symmetrical feature reconstruction in UNet^50^ shape and the 8^th^ stage performs fine feature fusion at the original scale. The upsampling and downsampling operations are symmetrical in layout and operated before the feature maps enter a certain transformer stage; only the spatial size of each video frame is changed while keeping the temporal length unchanged. Each transformer stage contains a stack of transformer blocks to jointly extract features, perform motion alignment between frames, and fuse the interframe information. The number of transformer blocks in the 8th stage is usually more than that in the 1^st^∼7^th^ stages (M>N). Finally, the feature maps are isotropically reconstructed along the simulated degradation axis (X/Y) to generate an isotropic subvolume. Global residual connections and skip connections are adopted to better reuse features and facilitate network learning.

**Fig. 1.**
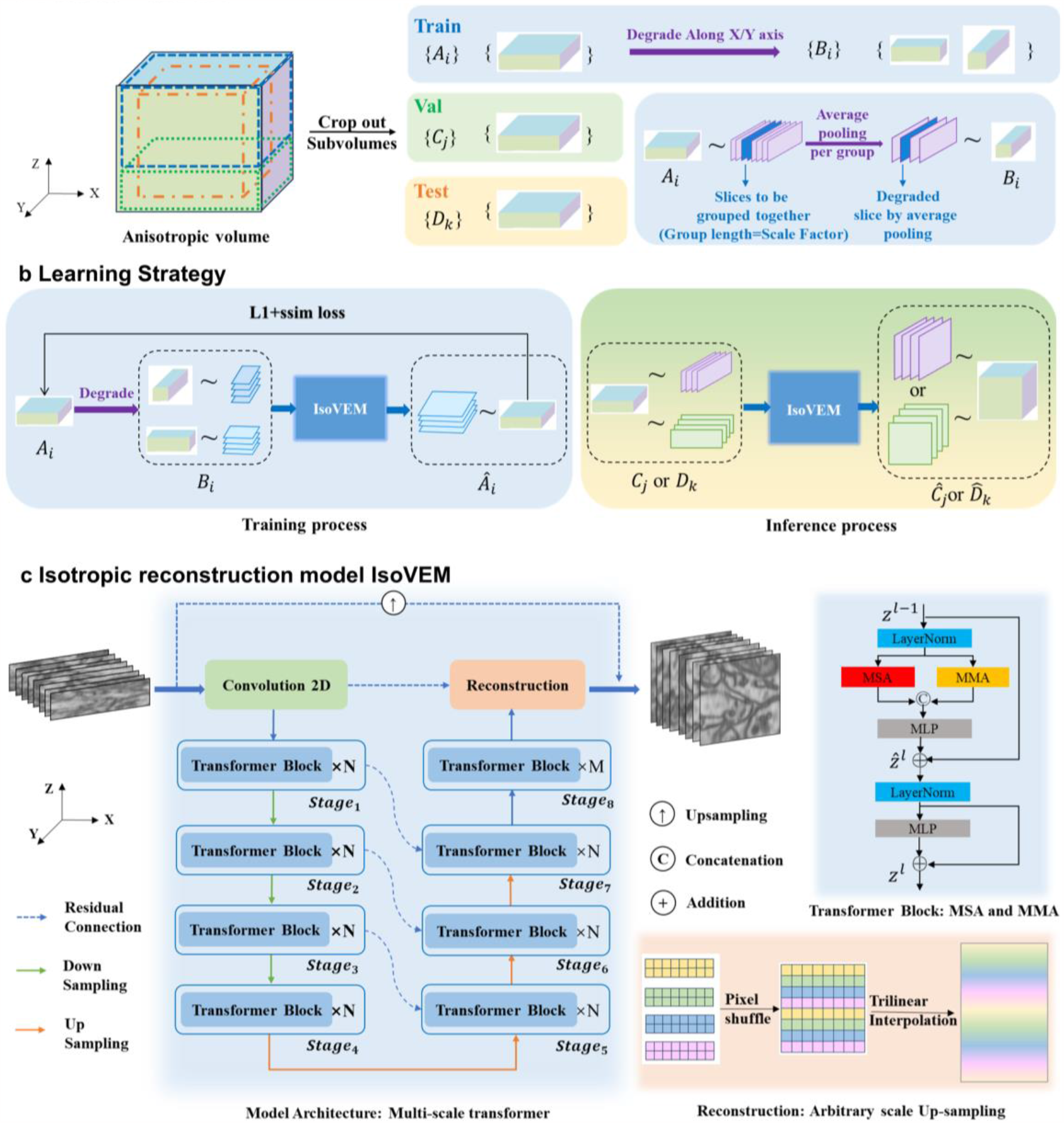
Algorithm design and process framework of IsoVEM. **a**, Generating subvolume sets for deep learning from raw anisotropic vEM volumes. The subvolumes are degraded along the X or Y axis to generate paired LR-HR data for self-supervised training using anisotropic average pooling as a degradation simulation method. *A*_*i*_, *C*_*j*_, *and D*_*k*_ represent the training set, validation set and test set, respectively; *B*_*i*_ represents the degradation volume along the X or Y axis. **b**, In the training and inference process, the model regards the input subvolume as a video clip along the Z-axis and X/Y-axis, respectively. The loss function is set as the weighted sum of the L1 and SSIM losses. *C*_*j*_ *or D*_*k*_ are the input data that need to be inferred by the IsoVEM network. *Â*_*i*_ is the output of the IsoVEM network, which is superresolved along the degradation axis. *Ĉ*_*j*_ or 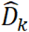 is the inferred volume. **c**, The model architecture is designed as a multiscale video transformer and upsampling reconstruction module. The 1^st^ to 7^th^ stages are constructed for multiscale feature extraction, and each scale has N transformer blocks containing MSA and MMA to perform implicit motion alignment. The 8^th^ stage has M transformer blocks designed for feature fusion. Arbitrary anisotropic restoration is achieved by trilinear interpolation after pixel shuffling inside the network.

Inspired by a video restoration transformer (VRT) ^51^, the transformer block includes two types of attention modules, MSA (multihead self-attention) and MMA (multihead mutual attention), to perform feature extraction and motion alignment (Supplementary Note 2). Self-attention^52^ (Extended Data Fig. 1a) uses self-similarity prior to images to make the feature more focused on important regions; this approach has been proven to be effective in various computer vision tasks.^53^ As shown in Extended Data Fig. 2, the self-attention map shows which areas of the input video clip are focused on, illustrating the long-distance dependency modeling ability across frames and the robustness to different structures. In contrast to self-attention, mutual attention (Extended Data Fig. 1b) emphasizes joint motion estimation and implicit feature alignment between two adjacent frames. The mutual attention map reflects the position correspondence between these two frames. Compared to explicit motion estimation and compensation methods such as optical flow^54,55^ and image warping, mutual attention has a simpler implementation, is more robust to large structural deformations between frames, avoids mismatch artifacts, and is more flexible for information warping in latent space. The multihead strategy^52^ is used for the above two attention mechanisms to enhance the semantic representation capabilities. Dividing and shifting attention windows (Supplementary Note 2) in the adjacent network layer to preserve long-distance dependency^56^ is necessary to reduce the computational cost of video processing, as shown in Extended Data Fig. 1c.

After the multiscale transformer stages, the feature maps are isotropically resolved by a reconstruction module composed of an anisotropic pixel shuffling^57^and trilinear interpolation to realize arbitrary-scale upsampling. First, the pixel shuffling operation is anisotropic, enhancing the resolution along only a single spatial axis direction. Second, the following trilinear interpolation enables plug-and-play arbitrary-scale upsampling, even for fractional scale factors. It should be noted that an anisotropic pixel shuffle can upsample at any positive integer scale factor, as long as the number of channels increases linearly with the scale factor. However, the scale factor during model inference must be consistent with that during the training phase. By following trilinear interpolation after an anisotropic pixel shuffle, the model can infer a scale factor that is inconsistent with the training phase, facilitating the transfer of the pretrained model to other vEM data with different anisotropic scale factors, even for fractional numbers. In addition, trilinear interpolation is effective for backpropagation and is lightweight without increasing the number of parameters or computations.

In summary, IsoVEM integrates feature extraction, motion alignment, and information fusion at multiple resolutions via multiscale video transformer stages and performs arbitrary-scale upsampling via the combination of anisotropic pixel shuffling and trilinear interpolation. Compared with existing isotropic reconstruction methods for vEMs, IsoVEM can trace structural movement between frames and maintain long-distance continuity, while 2D deep learning models cannot perform well. It also avoids the training instability and high-frequency artifacts that often occur in GAN-based models such as the SRGAN^29^ and CycleGAN^30^. It achieves excellent performance far beyond bicubic interpolation and close to supervised method^25^ (Fig. 2).

**Fig. 2.**
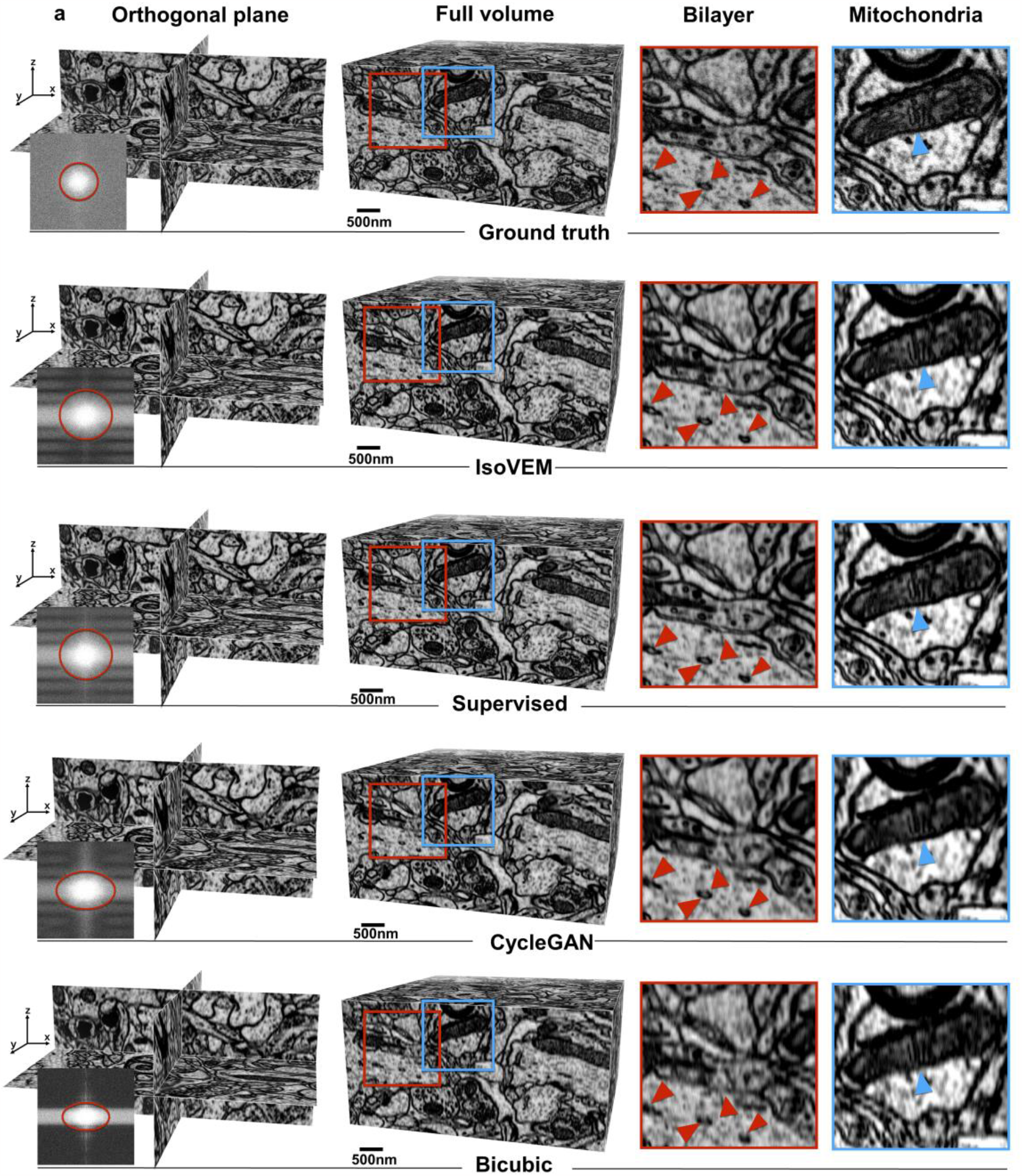
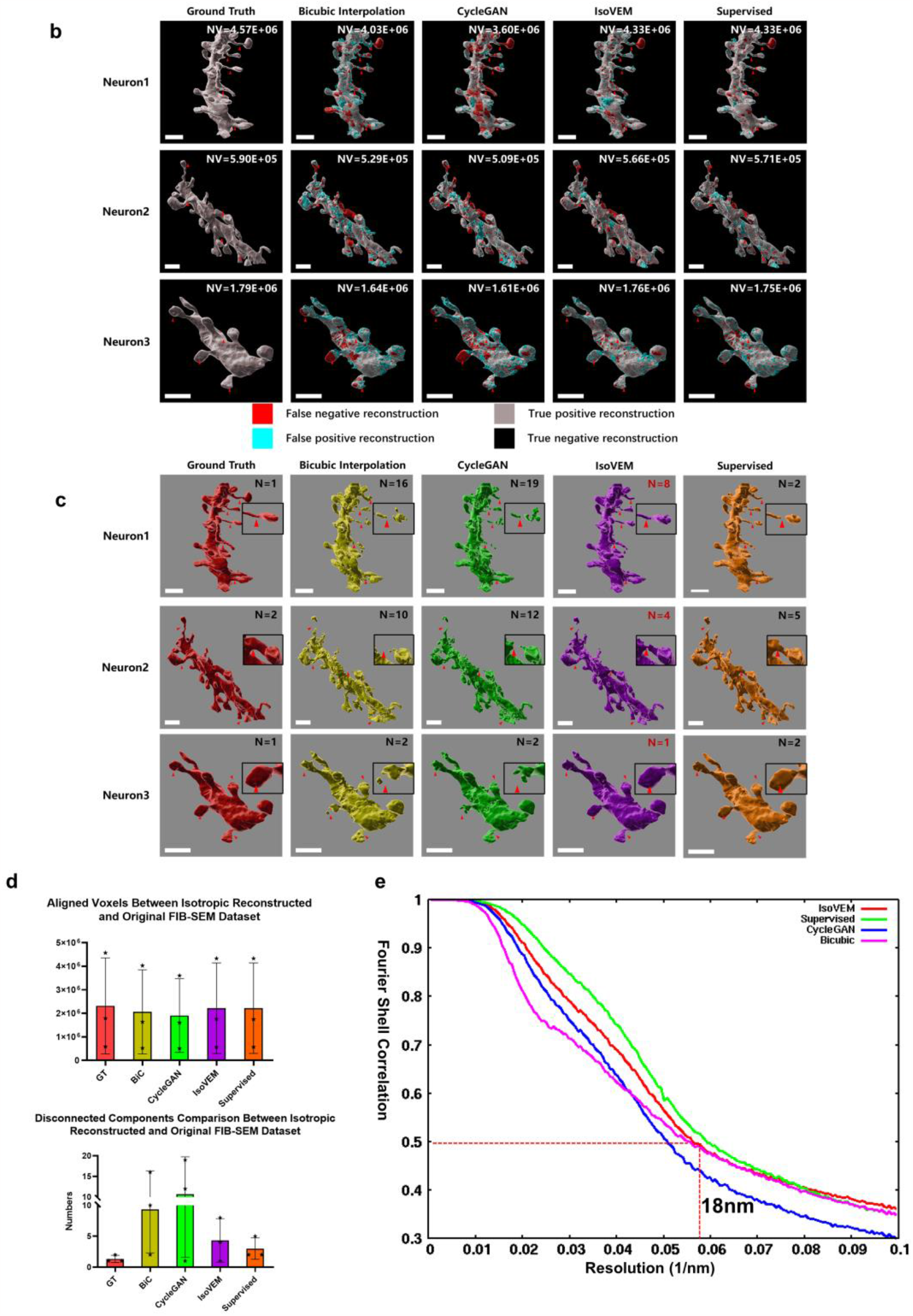
Comparison of isotropic reconstruction results of IsoVEM with existing methods on the EPFL dataset with the ground truth. **a**, The lines from top to bottom represent the ground truth, the reconstruction of IsoVEM, CycleGAN and bicubic interpolation, respectively. All methods are shown with an orthogonal view and full volume, while the power spectrum of the orthogonal x-z plane is presented. The local membrane structures and mitochondria from the full volume are highlighted with enlarged images and arrows (red arrows for membrane structures and blue arrows for mitochondria). **b**, Three neurons from the results of different reconstruction methods were segmented. Colocalization analysis between reconstructed neurons generated by different methods and ground-truth neurons was carried out. The values in the figure represent the volume of the colocalization in cubic microns. Except for the supervised method, the neurons segmented from the IsoVEM reconstruction result had the largest colocalization volume with the ground truth. **c**, Analysis of neuronal connectivity. The values in the figure represent neuron connectivity. A larger value indicates more breakpoints and poorer connectivity. Among the neurons isolated from 4 different isotropic reconstruction results, those extracted via the interpolation method exhibited the worst neuronal connectivity. The neuron connectivity of our IsoVEM method was better than that of CycleGAN. It was even better than the supervised method for neurons 2 and 3. **d**, The volume of overlapping regions and the number of discontinuous breaks of neurons in Fig. 2b and c were statistically analyzed. As Fig. 2b and Fig. 2c show, the bicubic interpolation method has the most breakpoints, the IsoVEM method was superior to the CycleGAN method, and the number of breakpoints for neurons from the ground truth was the lowest. The colocalization overlap volume of neurons for the interpolation method was the smallest, while the colocalization volume for IsoVEM reached the same level as that for the supervised methods. **e**, FSC cross-correlation calculation between the volume of each isotropic reconstruction method and the ground-truth volume. With the FSC 0.5 standard criterion and under the condition of a 5 nm imaging pixel in the x-y plane, IsoVEM recovered reliable axial information at 18 nm. Moreover, the FSC curve for IsoVEM (red curve) had a significantly greater correlation with the ground-truth structure, second to only the FSC curve of the supervised approach (green curve).

### 2.2. Isotropic reconstruction for the simulation dataset with ground-truth data

We downloaded isotropic data from mouse brain neurons collected by FIB-SEM from the EPFL website (https://www.epfl.ch/labs/cvlab/data/data-em/) and simulated 8 × anisotropy by averaging multiple frames every 8 sections. The sample contains not only subcellular structures, such as neurons and mitochondria, but also the ultrastructures of vesicles and cell boundaries at the nanometer level. This sample is typical for ultrastructure research using vEM technology, so we used these data to simulate serial section generation and test the effectiveness of the IsoVEM algorithm.

Compared with the supervised method^25^ and CycleGAN^27^, we found that the IsoVEM yields the most isotropic information in both the image space and Fourier space (Fig. 2a). The power spectra of the x-z planes show that IsoVEM can restore Fourier space information within the same diameter as the supervised network, while both CycleGAN^27^ and bicubic interpolation present elliptical diffraction information in Fourier space (Fig. 2a). More intuitively, the reconstruction results of the x-z planes of the IsoVEM method showed the same details as those of the supervised method ((Fig. 2a), and the resulting is significantly better than the bicubic interpolated one (Fig. 2a and Extended Data Movie 1). And more over, even the details were the same as those of the ground truth (Fig. 2a), such as the bilateral membranes (red box) and the mitochondrial cristae (blue box) (Fig. 2a). However, although the reconstruction of CycleGAN and bicubic interpolation also restore some axial information, the overall contrast is blurred, and structural details such as bilateral membranes cannot be presented.

We further segmented and extracted mouse neurons and mitochondria and performed morphological comparative statistics to confirm the effectiveness of IsoVEM further (Fig. 2b, c and d). For the sake of generality, we segmented three different locations of neurons in the reconstruction results of each method. The neurons recovered by IsoVEM had no jagged edges, and the voxel coincidence was closest to the ground truth (Fig. 2b and d). In addition to supervised methods, the segmentation breakpoints of neurons in the IsoVEM reconstruction results were also the smallest among all the deep learning-based methods and interpolation methods (Fig. 2c and d). For neurons 2 and 3, the number of broken points was even less than that of the supervised method. We believe that this is because IsoVEM makes full use of the 3D spatial information of the data through the video transformer model, so the reconstructed results were more continuous both in the whole volume and in the local ultrastructure, such as synapses. Mitochondrial segmentation via IsoVEM reconstruction revealed easily recognizable and complete mitochondria. Compared with those of the CycleGAN or interpolation methods, the mitochondria in the IsoVEM reconstructed results had the highest colocalization volume with the isotropic ground truth according to FIB (33.76 μm3), which was almost the same as that of the supervision method (33.87 μm3), while the interpolation method had the smallest colocalization volume (33.19 μm3) (Extended Data Fig. 3).

**Fig. 3:**
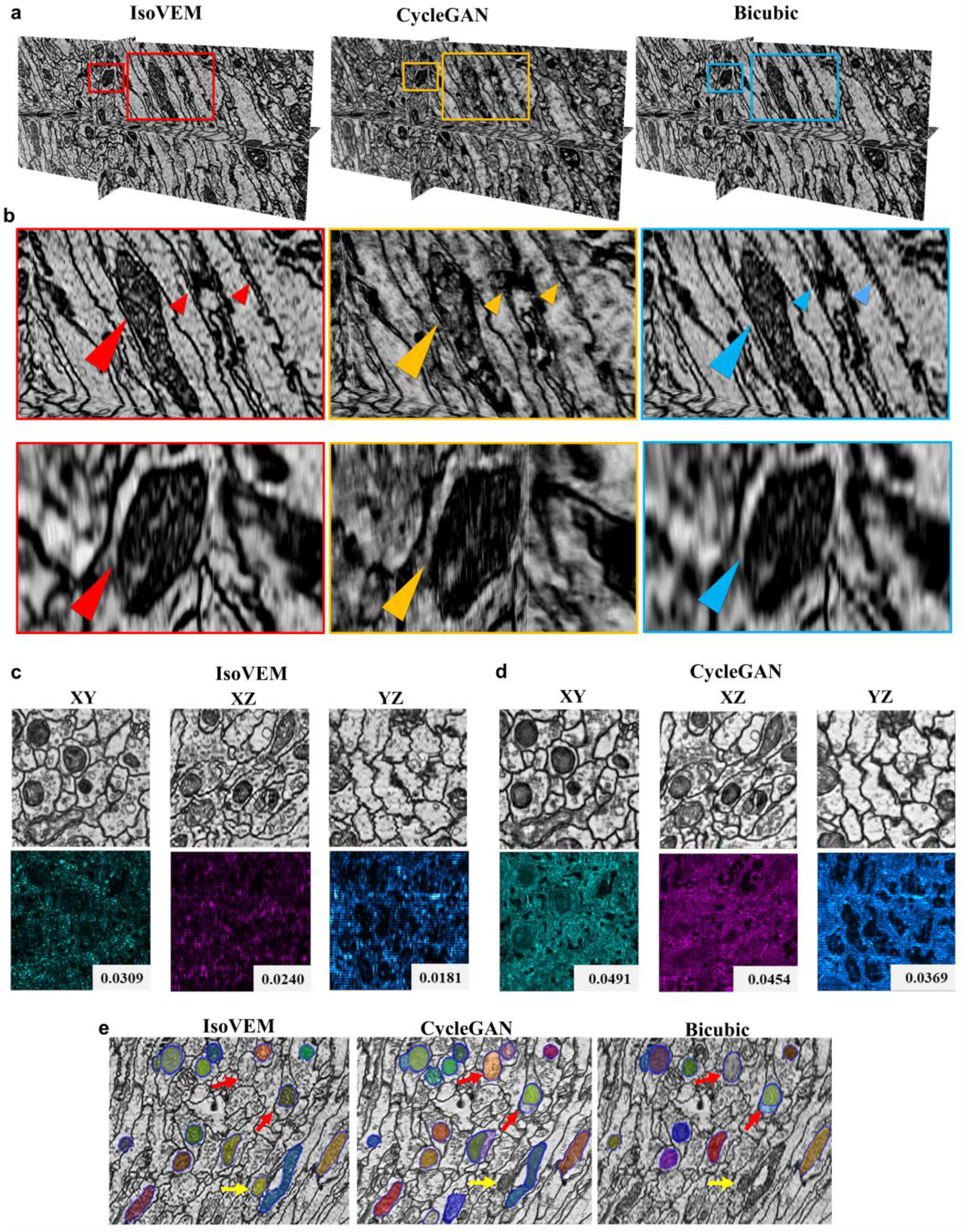
Comparison of real ssTEM data CREMI with an anisotropy factor of 10. **a**, The IsoVEM is clearer, and the sample details are more natural, while bicubic interpolation produces blurred structures, while CycleGAN exhibits layered artifacts. **b**, Highlighted regions for the above three reconstruction methods in a. IsoVEM (red rectangle) shows more complete and clearer structures at the boundaries of the membranous structure and the structure of the internal ridges of the mitochondria than CycleGAN (yellow rectangle) and bicubic interpolation (blue rectangle). **c, d**, Uncertainty map comparison for IsoVEM and CycleGAN. The artifact signal of CycleGAN was significantly larger than that of IsoVEM in three orthogonal directions, which indicates that the predictions of 8 kinds of rotation augmentation had larger standard deviations. IsoVEM reconstructed more stable and reliable structural information than CycleGAN. **e**, Mitochondria segmentation using SAM for reconstruction results of IsoVEM, CycleGAN and bicubic methods. Mitochondrial segmentation was performed using the instance segmentation method in the SAM.

FSC has been widely used as the gold standard for reliable reconstruction resolution estimation via cryoEM. Here, we introduce FSC calculations into these EPFL data with ground truth data. The FSC of the isotropic reconstruction result was calculated by comparing the Fourier shell cross-correlation with the ground-truth volume. Except for the supervised deep learning method, the FSC performance of IsoVEM is significantly better than that of the bicubic interpolation and CycleGAN methods (Fig. 2e). According to the FSC-0.5 criterion, IsoVEM could reach 18 nm isotropic resolution. These open EPFL data has 5 nm isotropic resolution, and we simulated the axial degradation with anisotropy 8, which means that the section thickness was simulated as 40 nm. The 18 nm recovery data imply that if people want to obtain a 20 nm axial resolution, slicing a sample with a 40 nm thickness will completely replace the 20 nm slicing strategy. This approach will save half the number of sections and further save at least 50% of the time during the entire process, such as sample preparation, 2D registration and 3D reconstruction. Moreover, electron microscopy and computer expenditure costs are reduced.

We also calculated the IsoVEM performance with different anisotropy factors and quantified the restoration quality by the pixel-based quantitative indicators PSNR, the structural similarity indices SSIM and MS-SSIM, and the visual perception indicator LPIPS for both the whole 3D reconstruction volume and three orthogonal planes (XY, XZ, and YZ). With an anisotropy factor of 4 or 8, all the above metrics for the IsoVEM were better than those of the methods based on bicubic interpolation or the CycleGAN (Extended Data Fig. 5 and Supplementary Table 2). This finding is consistent with the fact that the ultrastructures of neurons (Fig. 2) and mitochondria (Extended Data Fig. 3) observed by IsoVEM were closest to the ground truth. The double-layer membrane structure on the cell boundary region recovered by IsoVEM was even more continuous than that recovered by the supervised method (Fig. 2a, red arrows).

Taken together, the above findings demonstrate that IsoVEM can be used to recover ultrastructures with higher-resolution information in organelles and achieve reliable isotropic reconstruction results with higher FSC, lower reconstruction error and better visual perception (Fig. 2, Extended Data Figs. 3 and 4).

**Fig. 4.**
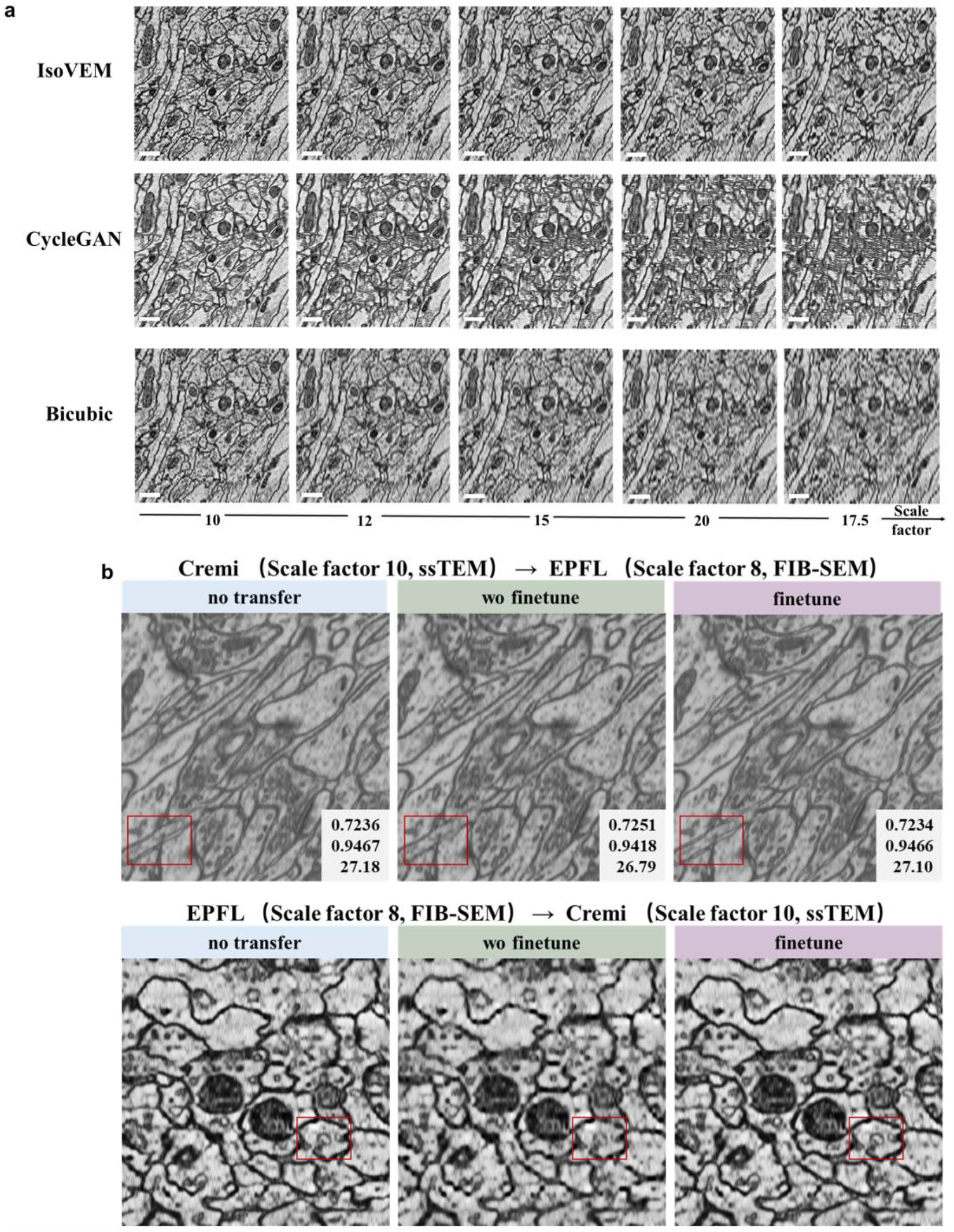
Model inference with different anisotropic factors and transfer robustness. **a**, We directly applied the IsoVEM trained on the 10x anisotropic CREMI dataset to simulate anisotropic EPFL data with arbitrary factors of 10x, 12x, 15x, 17.5x, and 20x. These factors covered odd numbers, integers, and decimals, ranging from 10x to fairly high 20x anisotropy. Compared with CycleGAN or bicubic interpolation, IsoVEM had clearer ultrastructural details for any anisotropic scale factor. **b**, After training on certain anisotropic factors, the IsoVEM easily performed network transfer learning between various anisotropic data. Here, when IsoVEM was trained using CREMI data with 10× anisotropy, the EPFL sample could be inferred and restored very well. The SSIM, MS-SSIM and PSNR for the transfer learning isotropic results were almost the same as those for the no transfer learning reconstruction. For the network trained by EPFL data with simulated 8x anisotropy, IsoVEM could also obtain CREMI structure details that were almost the same as those obtained without transfer learning. For both cases, the indicators or ultrastructure details for transfer learning after hyperparameter tuning were only slightly improved compared with the results without tuning. All scale bars are 500 nm.

**Fig. 5.**
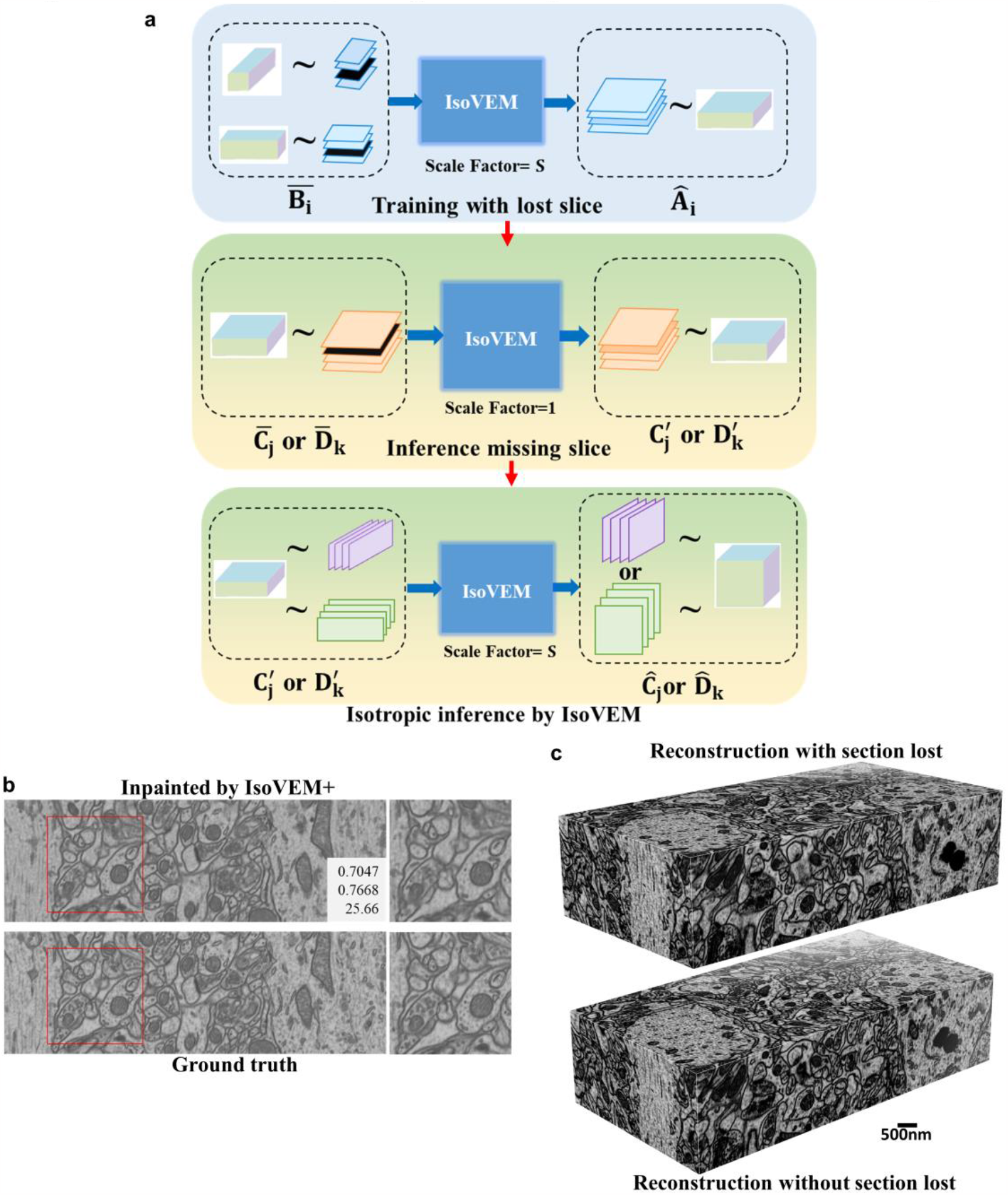
IsoVEM+ for simultaneous section inpainting and isotropic reconstruction. **a**, Pipeline for IsoVEM+ to jointly perform single-section inpainting and isotropic reconstruction on the condition of anisotropic data with defective sections. *Â*_*i*_,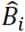, *Ĉ*_*j*_, and 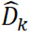 have the same meanings as in Fig. 1b.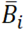 is the result of replacing the random portion of frames in 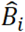 with a blank, and 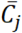, 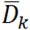 are the validation set and a test set with frames missing. 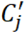, 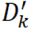 are the inpainting results of the validation set and a test set with frames missing, respectively. In the training stage, the model is trained jointly for single-frame inpainting and isotropic reconstruction, changing the input data from 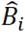 to 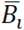, as described in Section 2.1. In the inference process, the model first performs section inpainting and generates 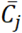, 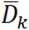 along the z-axis with an anisotropic scale factor of 1 and then performs isotropic reconstruction 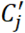, 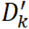 along the x/y axis based on 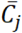, 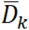. **b**, Verifying the model’s inpainting performance for lost frames on the EPFL dataset. The 11^th^ section of the EPFL data with 8× anisotropy simulation is removed, filled with zero, and then inpainted using IsoVEM+. The top image is the 11^th^ section inpainted with IsoVEM+, while the bottom image is the ground truth. **c**, 3D reconstruction of IsoVEM+ with section lost (top) and IsoVEM without section lost (bottom). Both have similar ultrastructural details.

### 2.3. Isotropic information recovery by IsoVEM for real data without ground truth

To further demonstrate the effectiveness of IsoVEM and its applicability to other vEM modal data, we tested the public ssTEM dataset Cremifrom website (http://cremi.org/). CREMI provides publicly available raw ssTEM images of the *Drosophila* melanogaster brain for algorithmic development and evaluation of neural circuits. These CREMI data were collected via ssTEM with an anisotropy factor of 10, and no ground truth was available for comparison. Therefore, we directly compared our method with traditional bicubic interpolation methods and CycleGAN networks. The results show that for such high-anisotropy data, IsoVEM reconstruction had the best structural details (Fig. 3a and b). The CycleGAN network recovered some axial information but could not recover the details of membrane-like structures, such as double layers of membranes (Fig. 3b). The results of the bicubic interpolation method were undoubtedly the worst; the overall axial structure was fuzzy, and the structural details were smoothed out (Fig. 3b). In contrast, the whole volume generated by IsoVEM provided continuous and natural structural information in 3D due to the space continuity learning advantage of video transformers in IsoVEM, while the volume obtained by CycleGAN or bicubic interpolation had obvious layer artifacts in the z axial direction. The reason is that CycleGAN is a two-dimensional model, and frame-by-frame inference along the x(y) axis generates layered discontinuous artifacts on the yz(xz) planes, which are difficult to eliminate even in the presence of training data augmentation. In contrast, IsoVEM keeps each section as a continuous video frame and adopts self-attention and mutual attention mechanisms for interframe motion alignment, thus maintaining the spatial continuity of the reconstructed volume.

To further evaluate the reliability of the model predictions on the CREMI dataset without the ground truth, artifact analysis was performed based on model uncertainty^33^. The training data were augmented based on 8 orthogonal rotations during model inference, so the model uncertainty could be measured by the standard deviation of those 8 predictions, and an uncertainty map of this deviation was generated to represent the reliability of each voxel position during model prediction. This is a disguised statistical verification method for the reality constraint when there is no truth value to compare. We compared the uncertainty maps of IsoVEM and CycleGAN on the CREMI dataset along each orthogonal axis (Fig. 3c) and found that CycleGAN had significantly more artifacts than IsoVEM, although they had similar structural textures. To verify the correlation between the uncertainty map and the error map, we evaluated the EPFL dataset with the ground truth (Extended Data Fig. 4). Both the uncertainty maps and error maps had the same distribution for bright regions, which appeared at membrane boundaries and complex structures. It should be noted that the defective area was removed by the network, so it was not reflected in the uncertainty map but still existed in the error map.

We attempted to segment the reconstruction result with the “segment anything model” (SAM)^58^. Compared with the CycleGAN results (Extended Data Fig. 6c and d) or bicubic interpolation (Extended Data Fig. 6e and f), the segmentation results for IsoVEM were obviously better (Extended Data Fig. 6a and b). The boundaries of the vesicles (Extended Data Fig. 6a) and membrane structures (Extended Data Fig. 6b) were clearly separated. For CycleGAN, due to the presence of artifacts, the edge segmentation of vesicles is not accurate, and the double-layer membrane is predicted to be a single-layer membrane. The mitochondrial segmentation of IsoVEM was also more accurate than that of the other methods, and the segmentation results for CycleGAN or the bicubic method showed that some mitochondria were not recognized (yellow arrow), while some areas that were not mitochondria were misidentified (red arrow) (Fig. 3e). This is because IsoVEM can reconstruct 3D space information continuously, especially when axial information is restored and staircase artifacts are removed. Therefore, the reconstruction of IsoVEM facilitates downstream analysis of ultrastructures and the study of biological mechanisms.

### 2.4. Transferability of IsoVEM

For the isotropic reconstruction of vEM, there are two problems associated with the scale factor and the model. First, different data have different anisotropy scale factors, which may be positive integers or decimals. The other is how to ensure that the model trained on the data with anisotropy a can be transferred to another dataset with anisotropy *b*(*b*≠*a*). We solved the above two problems through an arbitrary-scale upsampling module, which contains an anisotropic pixel shuffling step followed by trilinear interpolation (see Section 2.1 for details). This allows the IsoVEM to adapt to data with different anisotropic factors and make pretrained models transfer directly among them. We performed model inference on an arbitrary scale using the model trained on the 10x anisotropic CREMI dataset described in Section 2.3. For CycleGAN, to realize arbitrary-scale reconstruction during inference without changing the model weights, we simply changed the scale factor of the up- and downsampling module in the CycleGAN network.

Compared with the CycleGAN and bicubic interpolation methods, IsoVEM obtained complete and clearer ultrastructural information, and CycleGAN yielded obvious artifacts due to the mismatch between the network weight and anisotropy factor. Additionally, bicubic interpolation structures became blurred for large anisotropic factors (Fig. 4a). We also evaluate the EPFL dataset with ground-truth data (Extended Data Fig. 7) by simulating anisotropic data of any scale factor with an average pooling operation, as described in Section 2.1. Then, 5-nm isotropic data were reconstructed using different methods. The PSNR, SSIM, etc., validation indicators of these methods are shown in Supplementary Table 3. The IsoVEM obtained the best evaluation indicators among all three methods and different anisotropic factors.

The plug-and-play arbitrary-scale upsampling enables IsoVEM to transfer between different anisotropic data, including those collected using different electron microscopies, without the need for training on different data each time. For example, data collected via FIB-SEM were simulated as 8x anisotropic degradation along the z-axis, which could be recovered from the model trained on CREMI data collected via ssTEM with an anisotropy factor of 10 and vice versa (Fig. 4b and Supplementary Table 4). The performance of model fine-tuning across datasets was comparable to that of direct training and inference in the target domain data, which shows that the model has good transfer robustness (Fig. 4b and Supplementary Table 4). In this way, once the pretrained model is available, it can take less time to achieve good reconstruction through transfer learning in practical applications. (Fig. 4 b).

### 2.5. IsoVEM+: section inpainting and isotropic volume reconstruction simultaneously

At the sample preparation stage, artificial defects such as sample contamination, section wrinkles, broken sections, etc., are inevitable. For example, sections after diamond knife cutting may not be collected on tape, which results in missing sections during imaging. The occurrence of these phenomena brings further difficulties and challenges to 3D reconstruction of anisotropic vEM data. To solve these problems, we further developed the inpainting function for lost frames and defective frames, which is named IsoVEM+ and can achieve isotropic reconstruction and section inpainting simultaneously for defective data with missing sections, contamination, blurring, creases and so on.

In the first step, we used self-supervised training data {*B*_*i*_,*A*_*i*_} consistent with those in Section 2.1 but randomly replaced some input frames as blank frames 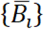 while keeping supervision {*A*_*i*_}unchanged. Then, IsoVEM training was performed as described in Section 2.1. This process involves simultaneous z-axis information estimation and x-y plane temporal inpainting for interframe filling. The second step removes defective sections by model inference. The defective area or lost frames of the input data are replaced with zero to generate 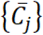 or 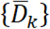, which represent the validation and testing sets, respectively, and a video clip is formed along the z-axis. The model performs inpainting only when the scale factor is set to 1 and generates 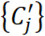 or 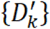, where contents are filled at blank defective frames based on contextual information. Then, the defective frames of the original data are replaced with the corresponding predictions of the model to generate a complete volume, which is used as input for isotropic reconstruction inference in the next step. The third step is basically the same as in Section 2.1 to perform isotropic reconstruction and obtain {*Ĉ*_*j*_}or. 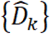 The defect removal data form a video clip along the z-axis, and the model performs isotropic resolution recovery only with a scale factor set to the anisotropy ratio.

We validated the performance of IsoVEM+ for single-defective section recovery and isotropic reconstruction on the EPFL dataset, which contains ground-truth data. The original data were simulated to be degraded by 8x anisotropy, after which twelve frames of the anisotropic volume were randomly selected and replaced with a blank along the z-axis. We trained IsoVEM+ and performed single-defective section recovery and isotropic reconstruction. Surprisingly, the frames that were lost in the simulation were recovered very well and had almost identical ultrastructures and information distributions to those of the corresponding frames in the ground truth (Fig. 5b and Extended Data Fig. 8). Through contextual feature training, the missing frame information is estimated accurately. The SSIM was greater than or near 0.6, and the PSNR was approximately 25 between the recovery frame and the ground truth for all the lost frames (Supplementary Table 5). This is another embodiment of the ability of IsoVEM to continuously recover spatial information. For isotropic recovery, we compared the results between the results with frame loss and without loss in Section 2.2, which involved degradation but without frame removal. We found that both the ultrastructure and the validation indices obtained with the ground truth were similar, while the SSIM and PSNR were slightly lower than those obtained without frame loss (Fig. 5c and Supplementary Table 5).

## 3. Discussion

The isotropic reconstruction method IsoVEM utilizes the 3D space continuity capability of video transformers to achieve outstanding reconstruction results and is suitable for the recovery of vEM data with any positive integer or decimal anisotropic factor. It can be applied to FIB-SEM, ssTEM, ssSEM and other different modal data and can be transferred from one to another, even in the case of different anisotropic factors. In addition, the model can handle the case of defective sections, which improves the robustness against disturbances from real imaging conditions such as contamination, missing sections, broken sections, and wrinkles. It provides reliable high-resolution isotropic data for downstream segmentation and ultrastructure analysis, such as neurons, mitochondria, etc.

IsoVEM can be further optimized and improved. First, the z-axis section degradation is approximated as average pooling in 3D, which can be replaced by other operations, such as frame extraction, or even by a deep learning network to simulate the degradation process. Second, IsoVEM achieves section inpainting while isotropic reconstruction. This attachable 2D section recovery function can be further expanded into section denoising or deblurring, simply by changing the input data during training accordingly. Third, in custom experiments, many intermediate steps of the IsoVEM, including the multiscale hierarchy, attention module, plug-and-play upsampling operator and loss functions, can be modified to achieve the best performance in their own experiments.

We verified that the pretrained IsoVEM model on a single dataset yields good transfer performance on other modal data. This indicates that IsoVEM pretrained on large-scale vEM data may become a universal vEM reconstruction model without the need for retraining. More tests for various datasets will be performed in future studies.

In addition, the image inpainting function in IsoVEM+ will, in turn, improve image registration accuracy. After reregistration, the images are then inpainted and isotropically reconstructed with IsoVEM+, which further improves the resolution of the results. Therefore, repeated iterations of IsoVEM+ and image registration will further improve the image quality of vEM reconstruction and bring many potential conveniences to the ultrastructure study of large-scale biological samples.

## 4. Method

### Image dataset preparation and preprocessing

The image datasets used in this work were prepared and preprocessed several times prior to analysis.

### Image dataset acquisition

The image volumes were obtained from two sources:

1. EPFL dataset: An isotropic FIB-SEM dataset^59^ of neural tissue from the hippocampal CA1 region was downloaded from the EPFL website (https://www.epfl.ch/labs/cvlab/data/data-em/). The original isotropic dataset represents a 5x5x5 μm^3^ volume at 5x5x5 nm/voxel resolution. This image was downsampled by 4x and 8x along the z-axis to generate simulated anisotropic datasets for validating isotropic reconstruction methods and comparative analysis.
2. CREMI dataset: Serial section TEM images of neural tissue were obtained from the CREMI repository. The datasets contain 193 sections at 4 × 4 × 40 nm/voxel resolution (Dataset A) (padded version, https://cremi.org/data/).

### Image registration

The “Register Virtual Stack Slices” plugin in ImageJ was utilized to align the serial tissue sections along the z-axis. This plugin aligns adjacent image sections using an automated feature matching algorithm based on the scale invariant feature transform (SIFT)^60-62^. For feature extraction, the “rigidly translated and rotated” model was used. For registration, the alignment models we used were “rigid - translate and rotate” and “moving least squares - maximal warping”. The optimal model was selected based on evaluation of the aligned image stack quality. When needed, elastic transforms were applied to refine the section alignment further. This plugin incrementally registers each section to the previous section to generate the aligned 3D image volume by extracting features between sections and finding optimal transforms to match them.

### Image cropping

The aligned image stacks were cropped down to specific regions of interest containing relevant ultrastructural tissue details. Cropping served two purposes: 1) focusing the analysis on relevant tissue areas and 2) removing large black areas introduced by rotation during image alignment. Cropping helped reduce file sizes and excluded unnecessary background regions from further analysis.

### Image enhancement

To enhance image contrast and standardize brightness across the image set, contrast limited adaptive histogram equalization (CLAHE) was applied^63^. This technique improves local contrast within small regions while limiting noise amplification. The CLAHE parameters were optimized to avoid overenhancement artifacts.

### Dataset preparation

The image datasets were extracted and preprocessed into three subsets: training, validation, and test sets. This enables model training, hyperparameter tuning, and unbiased evaluation.

The training dataset comprises the majority of images used to fit the model parameters. The full training datasets contain several blocks of anisotropic volumetric data (separation based on the spatial location of the defect sections). During training, random subvolumes of a fixed size (e.g., 16x128x128, 32x128x128) are sampled across these various blocks to form the training inputs foreach batch. This allows the model to train on diverse samples from a broader data distribution. To augment the training data, 8 orthogonal 3D rotations were applied.

A smaller portion of the stacks was clipped from the original complete blocks of anisotropic volumes for model validation. The validation subblocks are disjointed during the training process. The reconstruction of the validation data is completed at the best checkpoint for optimal isotropic quality. The validation subvolumes undergo identical preprocessing as the training data without augmentation.

The test dataset comprises one complete block of anisotropic EM volumes. Since training and inference occur on different dimensions, the same volumes used during training can also serve as test data. Using the same volumes enables direct evaluation of how well the model can superresolve in the z-dimension based on x-y training. The test volume also underwent standard preprocessing, similar to training data preparation, to eliminate reconstruction artifacts as much as possible.

### Network and hyperparameter design

For the model architecture, we set the number of stages of the multiscale transformer to 8: the 1st∼4th stage, the 4th∼7th stage, and the 8th stage for downsampling, upsampling and feature fusion, respectively. In the first 7 stages, each stage contains 4 (N=4) transformer blocks (TBs), in which 3 TBs contain both MMA and MSA followed by 1 TB containing only MSA. The 8th stage included 12TBs containing only MSAs. The channel sizes for the first 7 stages and the 8th stage are 40 and 60, respectively. For the attention module in the TB, we set the number of attention heads to 4 for both the MSA and MMA. The attention sizes are set as (2,2,16) and (2,2,20) for the EPFL and CREMI datasets, respectively. The larger the attention size is, the larger the model size. The model parameters of these two datasets are 1.4 M and 1.4 M, respectively.

In the model training stage, we set the training subvolume sizes to (16,16,128) and (16,16,160) for the EPFL and CREMI data, respectively. The batch sizes are all set to (2,2,1). We augment the input data by random cropping and 8 orthogonal 3D rotations. The loss function (Equation 3) we use is the weighted sum of L_1_(Equation 1) and the SSIM loss (Equation 2). The optimizer uses Adam^64^with β1 = 0.9 and β2 = 0.999. The learning rate is set to 1e^−3^, and an attenuation strategy is not used. It takes approximately 80k∼150k iterations to reach convergence.

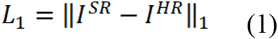

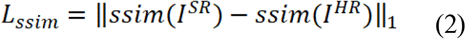

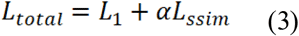

where *I*^*HR*^ is a high-resolution image and *I*^*SR*^ is a superresolution reconstructed image. The weight parameter *α* is set to 1 in all our experiments.

In the model inference stage, we generate the testing subvolumes by cropping the large volume with overlap (8,8,8) in 3D, the size of which is consistent with the training volume. Model inference on those subvolumes in an augmentation manner by 8 orthogonal 3D rotations. The generated isotropic subvolumes are then stitched in 3D to obtain the whole volume according to the cropping coordinates. Linear fusion is adopted for the overlapping area during stitching.

The code was implemented in Python 3.8 and PyTorch 1.8.1. Model training and testing were conducted on a single RTX 3090 NVIDIA GPU card for all the experiments. Depending on the size of the vEM dataset, it took approximately 0.8∼1.2 days to train the model and several hours for model inference.

### Evaluation metric

We used four quality assessment metrics to evaluate the isotropic reconstruction performance on the EPFL-simulated degraded data:

#### (1) SSIM, Structural Similarity Index Measure

The SSIM score^65^ can reflect the perception of the human visual system on local structural changes in images. The computation is based on three factors: luminance (*l*), contrast (*c*) and structure (*s*). The overall index is their multiplicative combination.

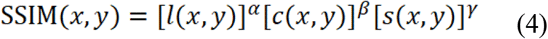

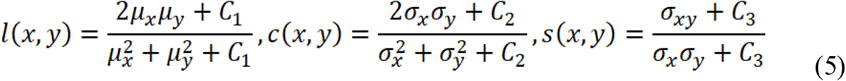

where `α=β`=*γ*=1 and C_3_=C_2_/2. *μ*_*x*_, *μ*_*y*_, *σ*_*x*_, *σ*_*y*_ and σ_xy_ are the local means, standard deviations, and cross-covariance for images *x* and *y*, respectively. The SSIM values range between 0 and 1; the closer the value is to 1, the better the performance.

#### (2) MS-SSIM, Multiscale Structural Similarity Index Measure

MS-SSIM^66^ extends the SSIM measurement to multiple scales, where the image at scale M is half the image at scale M-1. Through such adaptation, the MS-SSIM is more robust than the SSIM for various image resolutions.

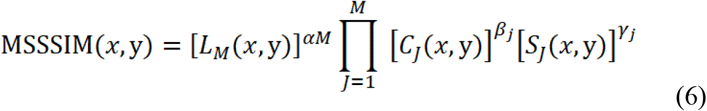

where *β*_1_= *γ*_1_ = 0.0448, *β*_2_= *γ*_2_ = 0.2856, *β*_3_= *γ*_3_= 0.3001, *β*_4_ = *γ*_4_ = 0.2363, and. *α*_5_ = *β*_5_ = *γ*_5_ = 0.1333.

#### (3) PSNR, peak signal-to-noise ratio

The PSNR is the ratio between the maximum possible signal power of an image and the power of corrupting noise that affects the image quality. In the actual calculation, we estimate the PSNR of an image by comparing it to an ideal clean image with the maximum possible power. A high PSNR indicates good image quality.

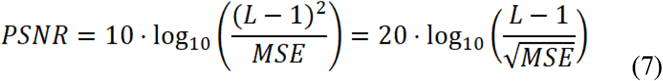

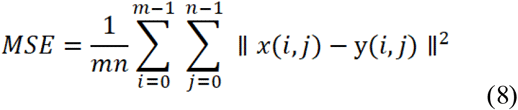

where L is the number of maximum possible intensity levels in an image. *x* and *y* are two images to be compared with the same height *n* and width *m*.

#### (4) LPIPS, learned perceptual image patch similarity

LPIPS^67^ calculates the perceptual similarity between two images and matches human perception well. LPIPS first generates activations of two 2D images by means of a pretrained network (such as VGGNet, which is pretrained on ImageNet), and then calculates the feature differences on different layers and spatial dimensions. Since the vEM data are 3D, we calculate the LPIPS of each frame and average it along a certain axis. A low LPIPS indicates that the image patches are perceptively similar.

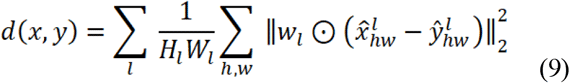

where 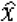 and *ŷ* denote the activation of two image patches. *l* represents the layer of the network, and *H* and *W* represent the height and width of the image, respectively.

#### (5) FSC, Fourier shell cross-correlation

FSC^68^ is a function of spatial frequency and was first introduced by Harauz and van Heel. The method calculates the normalized cross-correlation coefficient between two 3D volumes on corresponding shells with radius *r*_*i*_ in Fourier space. We calculated the FSC between the IsoVEM reconstructed volume and the corresponding ground-truth volume. The signal-to-noise ratio of vEM data is much better than that of cryoEM data. Therefore, we use the FSC-0.5 standard, which is more suitable for situations where the signal is much more significant than the noise ^68^.

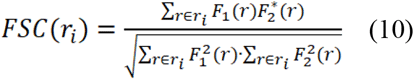

where *F*_1_(*r*) and *F*_2_(*r*) are the Fourier transforms of the IsoVEM reconstructed volume and ground-truth volume, respectively, at radius *r* in Fourier space; 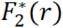 is the conjugator of *F*_2_(*r*)

### Evaluation of the Restored vEM Volumes Postprocessing

Reconstruction was performed on the complete datasets using the checkpoints with the best validation performance. A comprehensive evaluation was performed on the restored vEM volumes. The restored output was imported into MIB for segmentation. The GraghCut-based semiautomatic segmentation pipeline and deep learning-based automatic segmentation pipeline were used in the MIB to delineate critical ultrastructural boundaries^69,70^.

For the downsampled EPFL dataset, the neurons randomly selected from the ground truth were segmented by labeling the foreground and background features using the GraphCut tool. We used the same predefined features to segment the corresponding neurons in the restored datasets. The segmentation results of these volumes restored by various methods and ground truths were imported into Imaris for surface rendering. We compared the number of disconnected components and overlapping volume/voxels of the surface to assess the pixel similarity and topological relationship similarity between the reconstructed data and the original data. The F1 score was calculated as the harmonic mean of the precision and recall scores of a model. It is defined as follows:

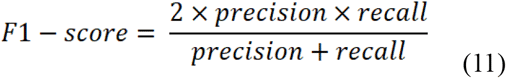

where precision is the number of true positive results divided by the number of all positive results, including those not identified correctly, and recall is the number of true positive results divided by the number of all samples that should have been identified as positive.

For the CREMI dataset, we used the segment anything model SAM (https://segment-anything.com/) to assess the topological structural relationships of the datasets via segmentation of membranes, vesicles, mitochondria, etc.

### Statistical analysis

Surface-rendered Imaris models were used to characterize neurons and organelle morphologies in 3D. We used Excel to integrate and analyze the preliminary statistical data and GraphPad Prism 9.5 to complete the plotting and significance testing analyses.

## Data availability

1. The EPFL dataset was downloaded from the EPFL website (https://www.epfl.ch/labs/cvlab/data/data-em/).
2. The CRIMI dataset was obtained from the CREMI repository (https://cremi.org/data/).

## Code availability

The source codes of IsoVEM are available at https://github.com/cbmi-group/IsoVEM.

## Acknowledgments

This work was supported by the National Natural Science Foundation of China (322371248 to Y.Z., 31971289, 91954201 to G.Y., 92254306 and 31925026 to F.S.), the National Key Research and Development Program of China (2021YFF0704300 to Y.Z. and 2021YFA1301500 to F.S.), and the Strategic Priority Research Program of the Chinese Academy of Sciences (XDB37040402 to G.Y.). We thank Y. Feng and C. L. Liu from the Center for Biological Imaging (CBI), Institute of Biophysics, Chinese Academy of Science, for their help in image segmentation. We acknowledge the services provided by X. X. Li from CBI, Institute of Biophysics, Chinese Academy of Science for ssSEM data collection. We thank S. D. Yang from the Institute of Biophysics, Chinese Academy of Science, for his help with the data preprocessing and L. R. Li from the Institute of Automation, Chinese Academy of Science, for her help with the interpolation method survey.

## Author contributions

F. S. conceived the concept. Y.Z. supervised the study with support from F.S. and G.Y. J.H. proposed the model architecture and conducted all the model training and testing. J. H. conducted code replication of the supervised method and CycleGAN. W. H. S. performed preprocessing of the test data and segmentation of the EPFL data. J. H. performed segmentation of the CREMI data. Y.Z. analyzed all the data and results. Y.Z. wrote the manuscript with input from all the coauthors. Y.Z., G.Y. and F.S. acquired funding for this project.

## Notes

### Competing Interest Statement

The authors have declared no competing interest.

### Summary of Updates

The title was updated; The name of this algorithm was renamed from EMformer to IsoVEM; The word session in the whole article was updated as section.Authoer affiliations updated.

