## Extended Data Figure 1 for "IsoVEM: Isotropic Reconstruction for Volume Electron Microscopy Based on Transformer"

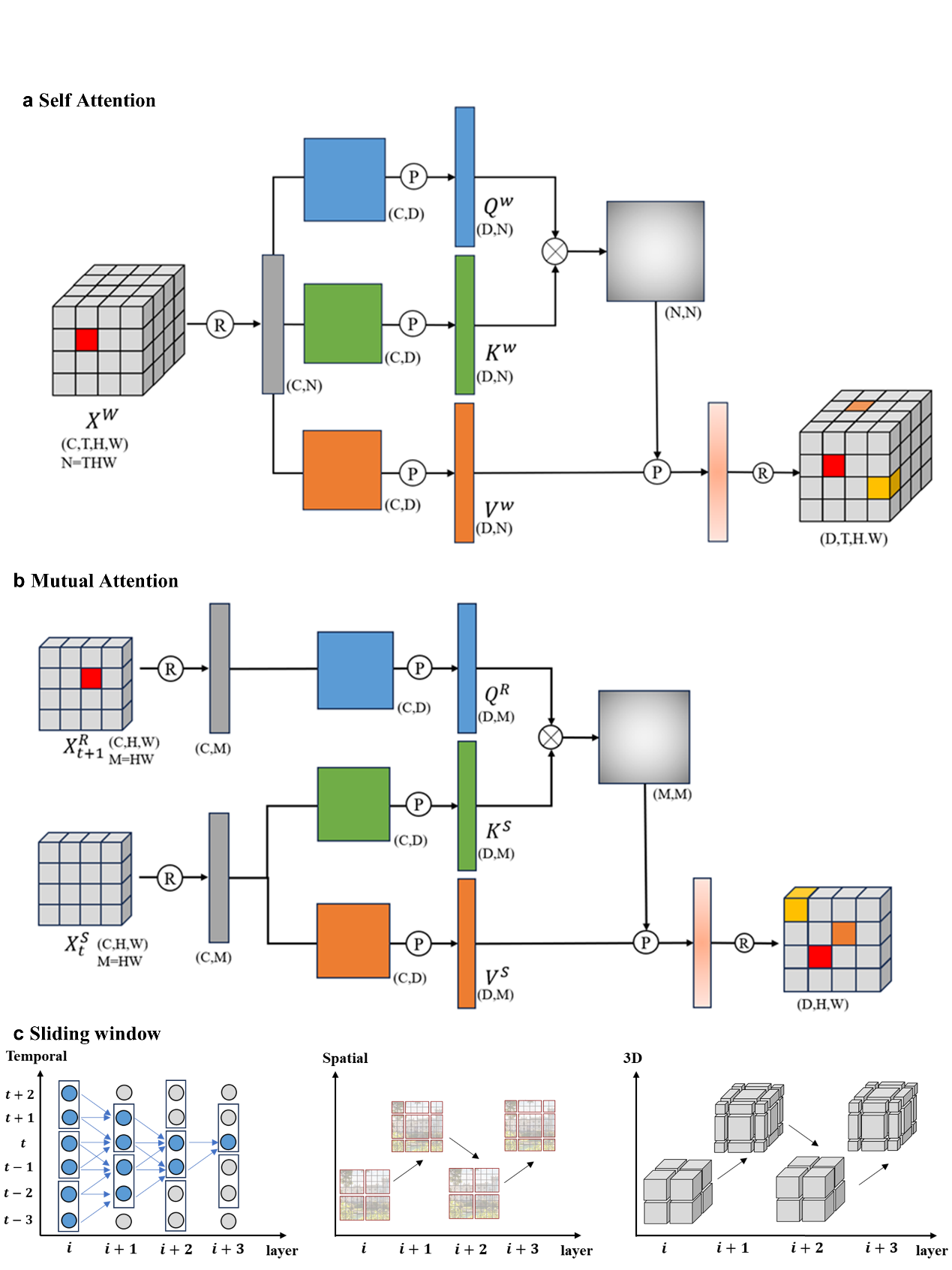
**Extended Data Fig. 1 | The attention mechanism in IsoVEM, considering the sub-volume latent feature as latent video clip.** **a,** Computation graph of self-attention based on input video clip itself. **b,** Computation graph of mutual attention based on two adjacent frames. c. Shifted window mechanism in temporal, spatial and 3D. The input data is partitioned into 3D window clips at each layer and shifted for every other layer to enable cross-clip interactions and reduce computation.
