## Extended Data Figure 2 for "IsoVEM: Isotropic Reconstruction for Volume Electron Microscopy Based on Transformer"

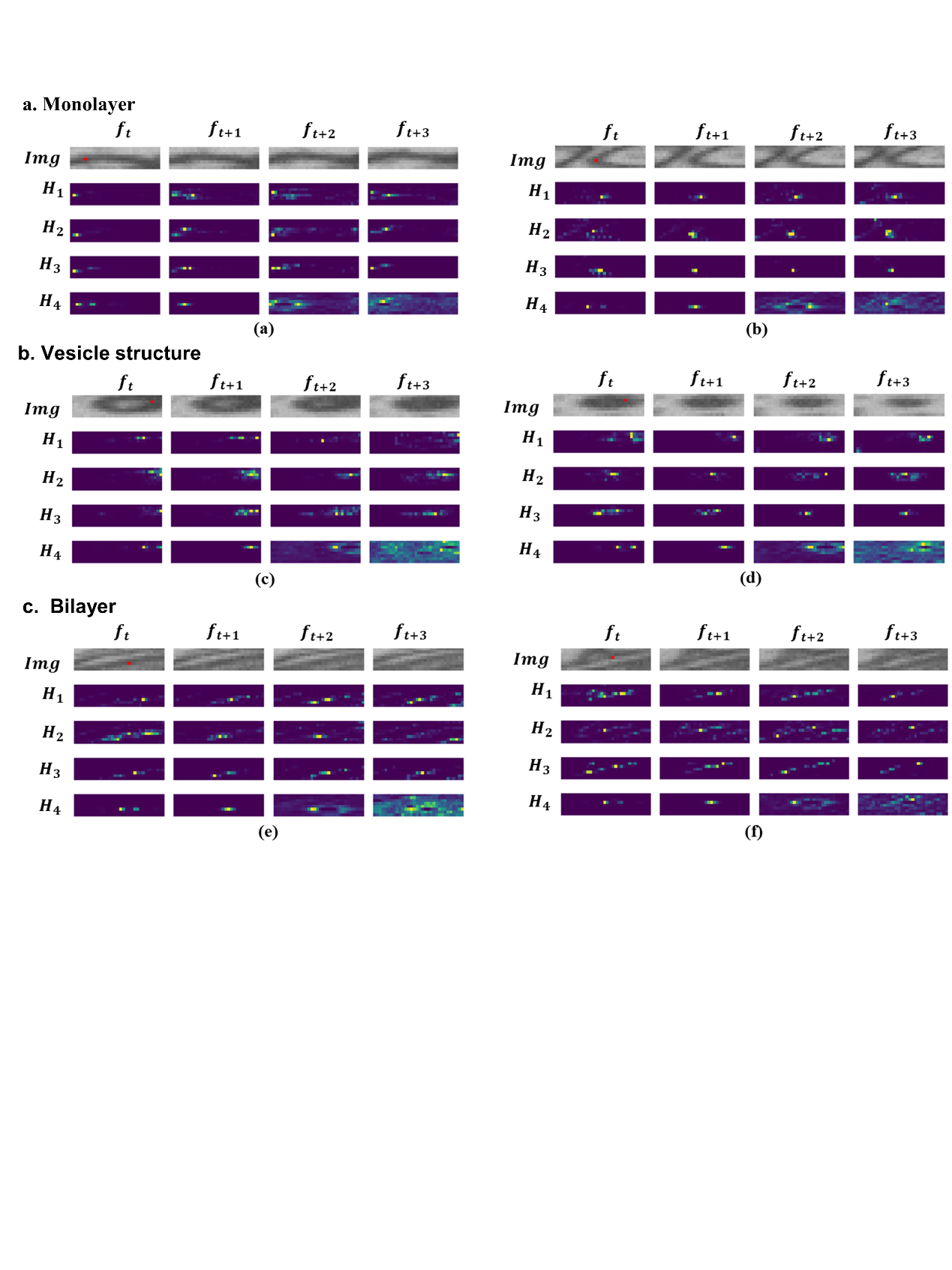

**Extended Data Fig. 2 | Visualization of attention map on different biological features on EPFL dataset.** The attention map is extracted from stage1 module of IsoVEM without down-sampling and window size is set to (4,8,32). In each subgraph, the first row represents the original image patches at the same position of different frames, while the remaining rows are the visualization of the attention weights of four different attention heads. Query pixels are marked by red dots in the first frame. For some classical structures in connectomics with biological ultrastructure such as mono-layer (**a**), vesicle structure (**b**) and bilayer (**c**), the attention map correctly focuses on specific areas similar to query sites, and the inter frame motion direction of the attention area in the attention map is basically consistent with the actual structure.
