## Extended Data Figure 3 for "IsoVEM: Isotropic Reconstruction for Volume Electron Microscopy Based on Transformer"

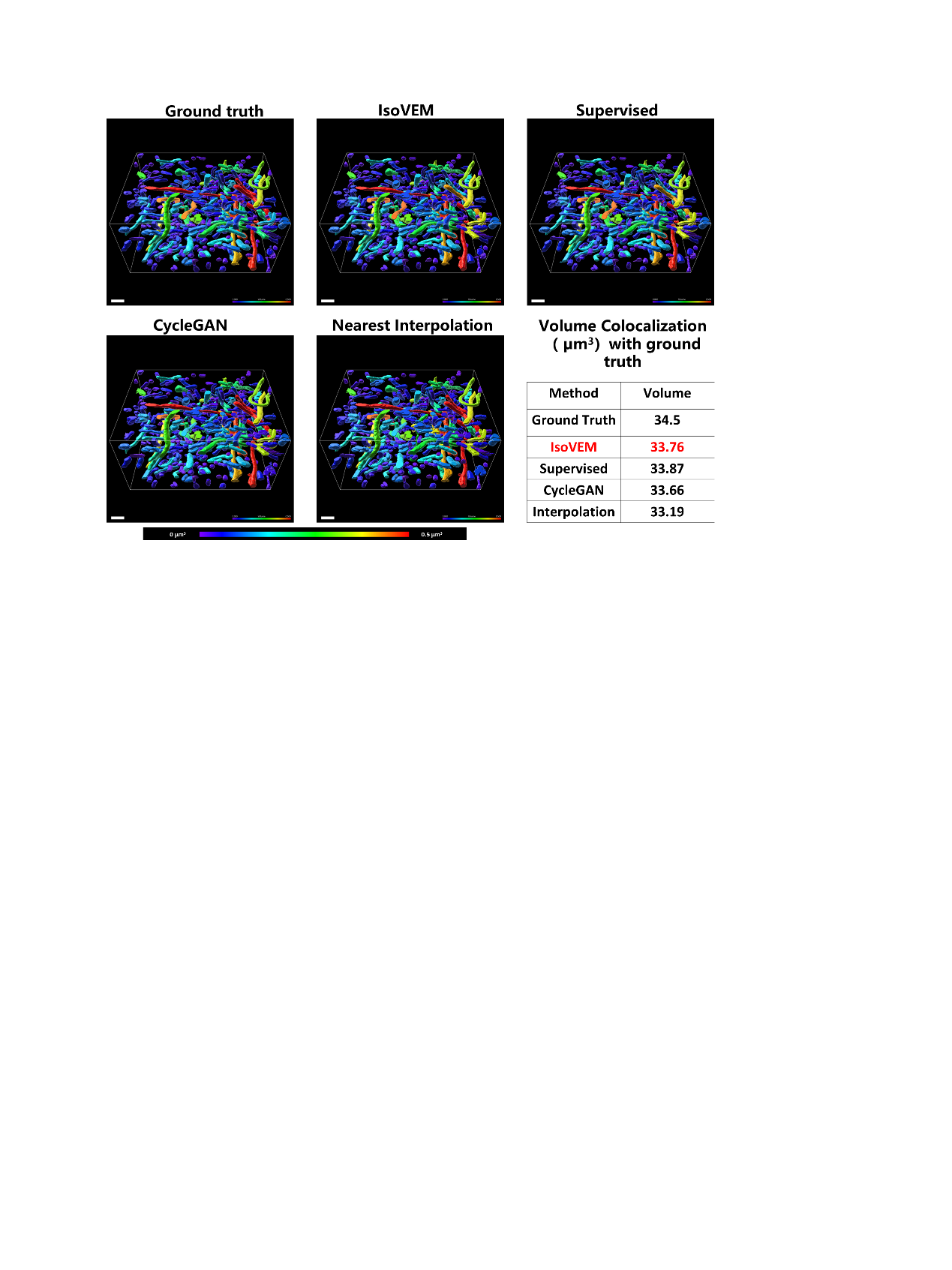

**Extended Data Fig. 3 |** **Mitochondria segmented from EPFL for IsoVEM and others isotropic reconstruction method.** Morphologically, there is no significant difference in mitochondria in the reconstructions of the various methods. However, except supervised method, the colocalization volumes between mitochondria reconstructions and ground truth show that IsoVEM is better than other methods. Scale is 1**µm.**
