## Extended Data Figure 4 for "IsoVEM: Isotropic Reconstruction for Volume Electron Microscopy Based on Transformer"

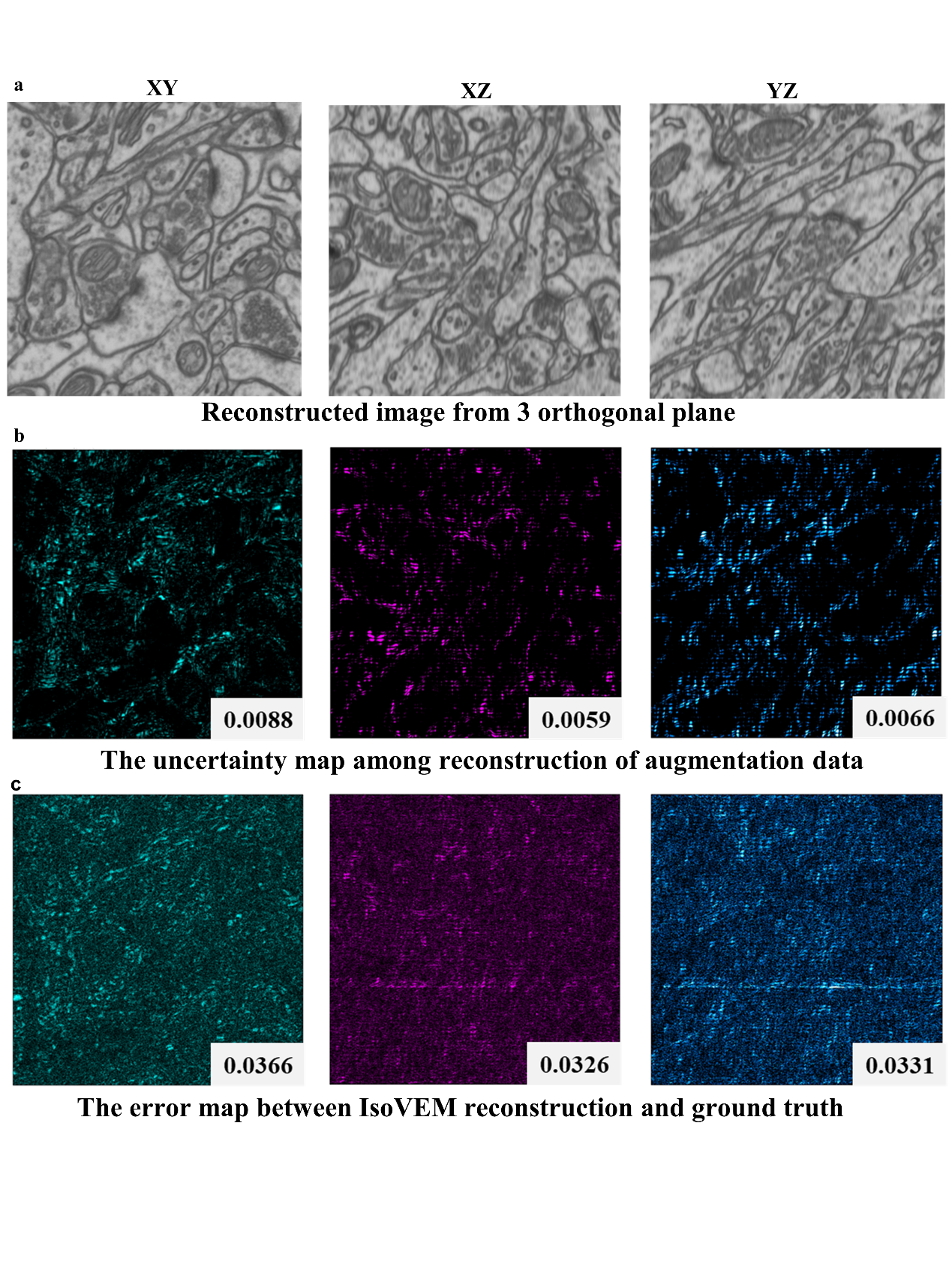

**Extended Data Fig. 4 | Verifying the correlation between uncertainy map and error map on the EPFL dataset.** **a,** Three images from orthogonal planes of IsoVEM reconstructed volume. **b,** Standard deviation of 8 reconstructed volumes obtained by 8 kinds of orthogonal rotation augmentation is used as uncertainty map. The indicator is the average value of uncertainty map calculated on the normalized image. **c,** The error map between IsoVEM reconstructed volume and isotropic ground truth. The distribution of bright areas in the uncertainty map and error map are basically consistent. The indicator is the average value of error map calculated on the normalized image. In addition, the volume reconstructed by IsoVEM automatically removes the defects from the original image, so the defects signal only exist in the error map and not in the uncertainty map.
