## Extended Data Figure 5 for "IsoVEM: Isotropic Reconstruction for Volume Electron Microscopy Based on Transformer"

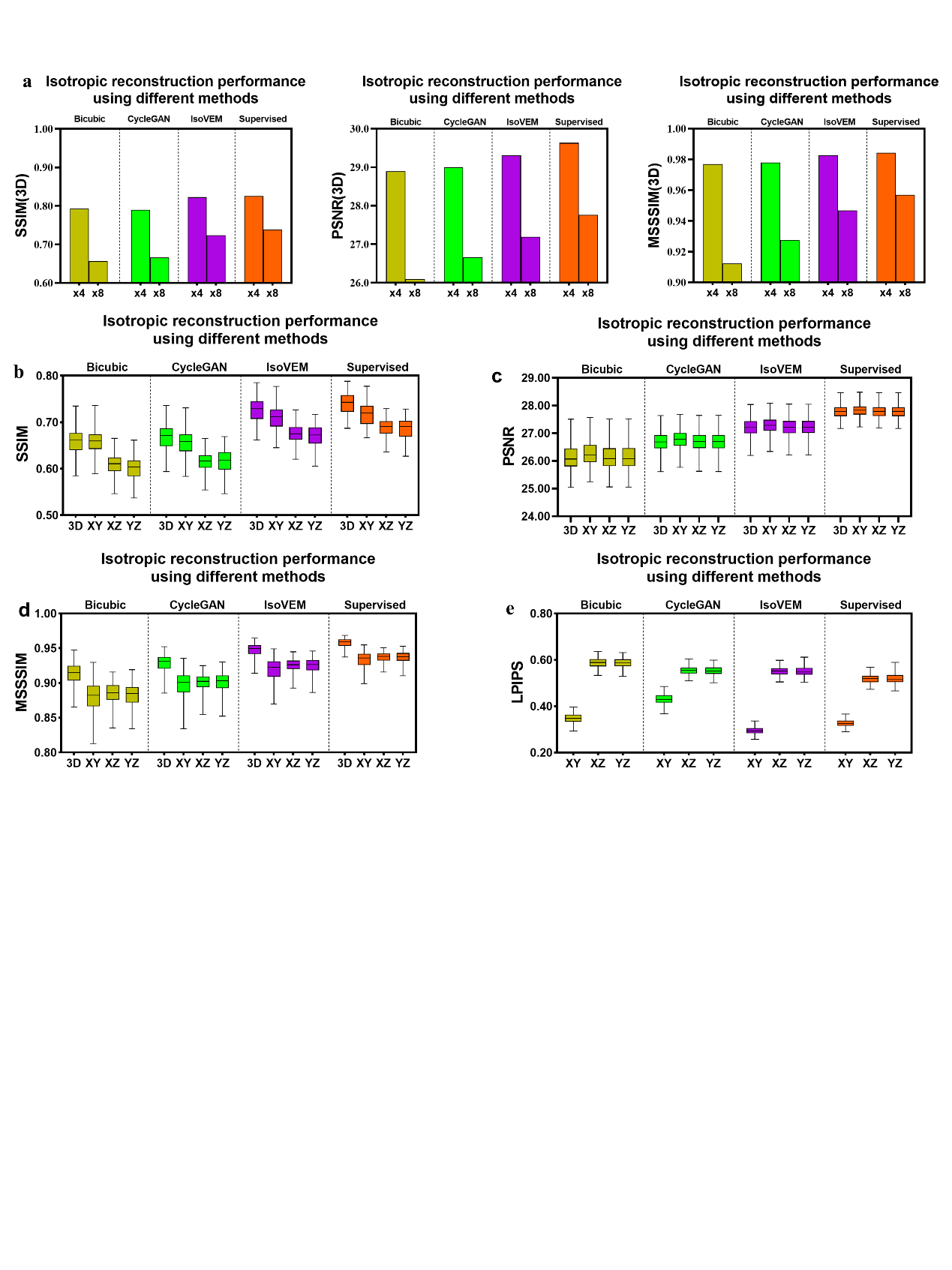

**Extended Data Fig. 5 | Comparison of evaluation metrics for IsoVEM and other methods.** **a,** The EPFL dataset was degraded along z axis by 4 and 8 times respectively and then reconstructed using each method. The SSIM, PSNR and MSSSIM were calculated between the reconstruction and ground truth. IsoVEM presents better indicator than bicubic interpolation or CycleGAN method, and closet to supervised method. **b,** SSIM of 3D reconstructed volume and XY, XZ, YZ orthogonal planes when different method performs 8x isotropic reconstruction. **c,** PSNR of 3D reconstructed volume and XY, XZ, YZ orthogonal planes when different method performs 8x isotropic reconstruction. **d,** MSSSIM of 3D reconstructed volume and XY, XZ, YZ orthogonal planes when different method performs 8x isotropic reconstruction. **e,** LPIPS of XY, XZ, YZ orthogonal planes when different method performs on 8x anisotropic restoration.
