## Extended Data Figure 6 for "IsoVEM: Isotropic Reconstruction for Volume Electron Microscopy Based on Transformer"

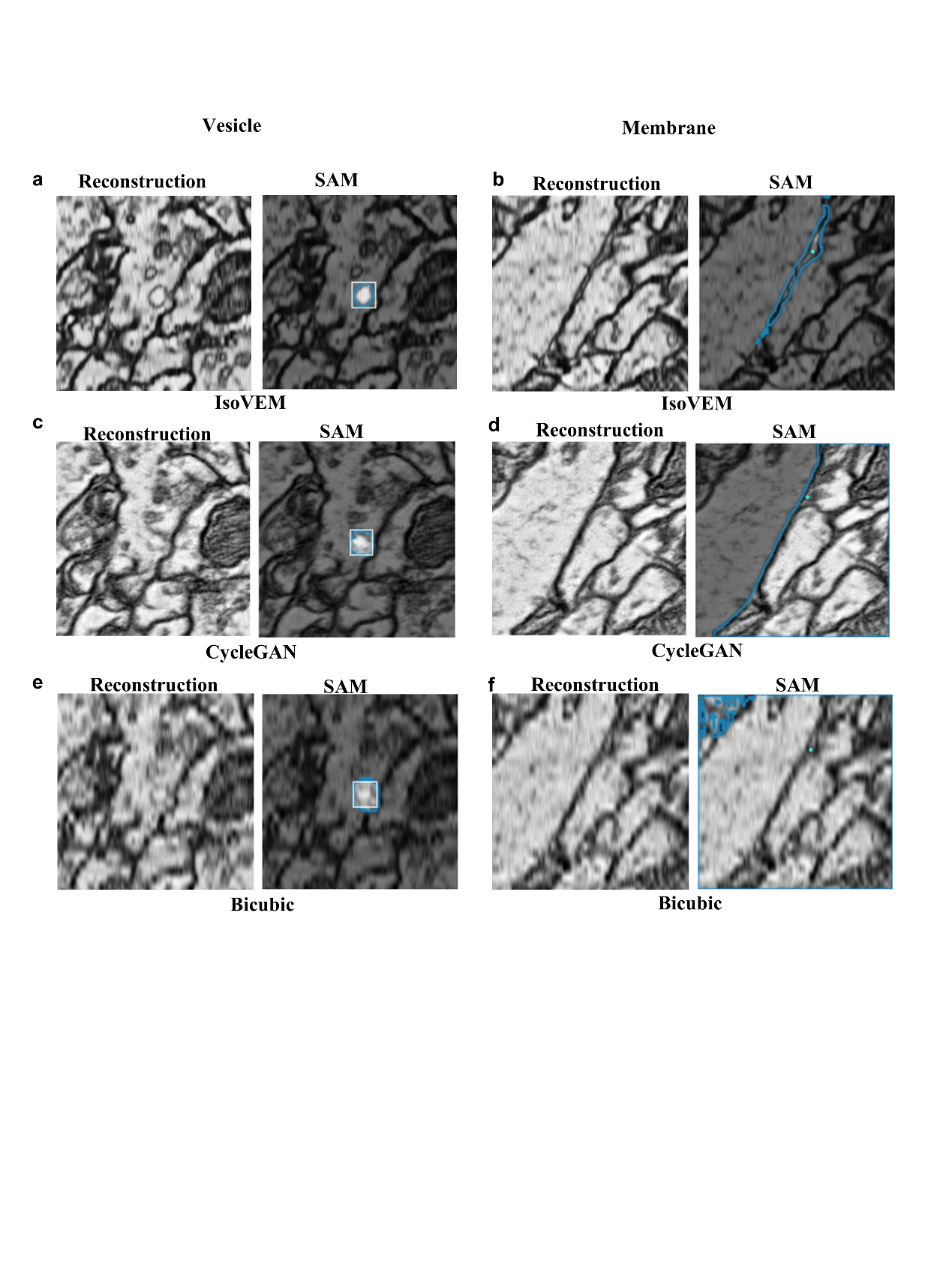

**Extended Data Fig. 6 | Using SAM to segment various structures in axial orthogonal plane (XZ) reconstructed by different methods for Cremi data. a,c,e,** Square prompt was used to segment vesicle of reconstruction results from IsoVEM, CycleGAN, bicubic interpolation respectively. IsoVEM's vesicle reconstruction has clear boundaries, while CycleGAN's boundaries are less complete, and boundaries in bicubic result are blurry. **b,d,f,** Point prompt was used to segment membrane of reconstruction results from IsoVEM, CycleGAN, bicubic interpolation method respectively. Double-layer membrane can be segmented out accurately for IsoVEM's reconstruction result. Although CycleGAN has good visual effect, it does not resolve those fine structures and the double-layer membrane cannot be recognized. Bicubic interpolation produces ambiguous segmentation of the whole image due to blurred edges.
