## Extended Data Figure 7 for "IsoVEM: Isotropic Reconstruction for Volume Electron Microscopy Based on Transformer"

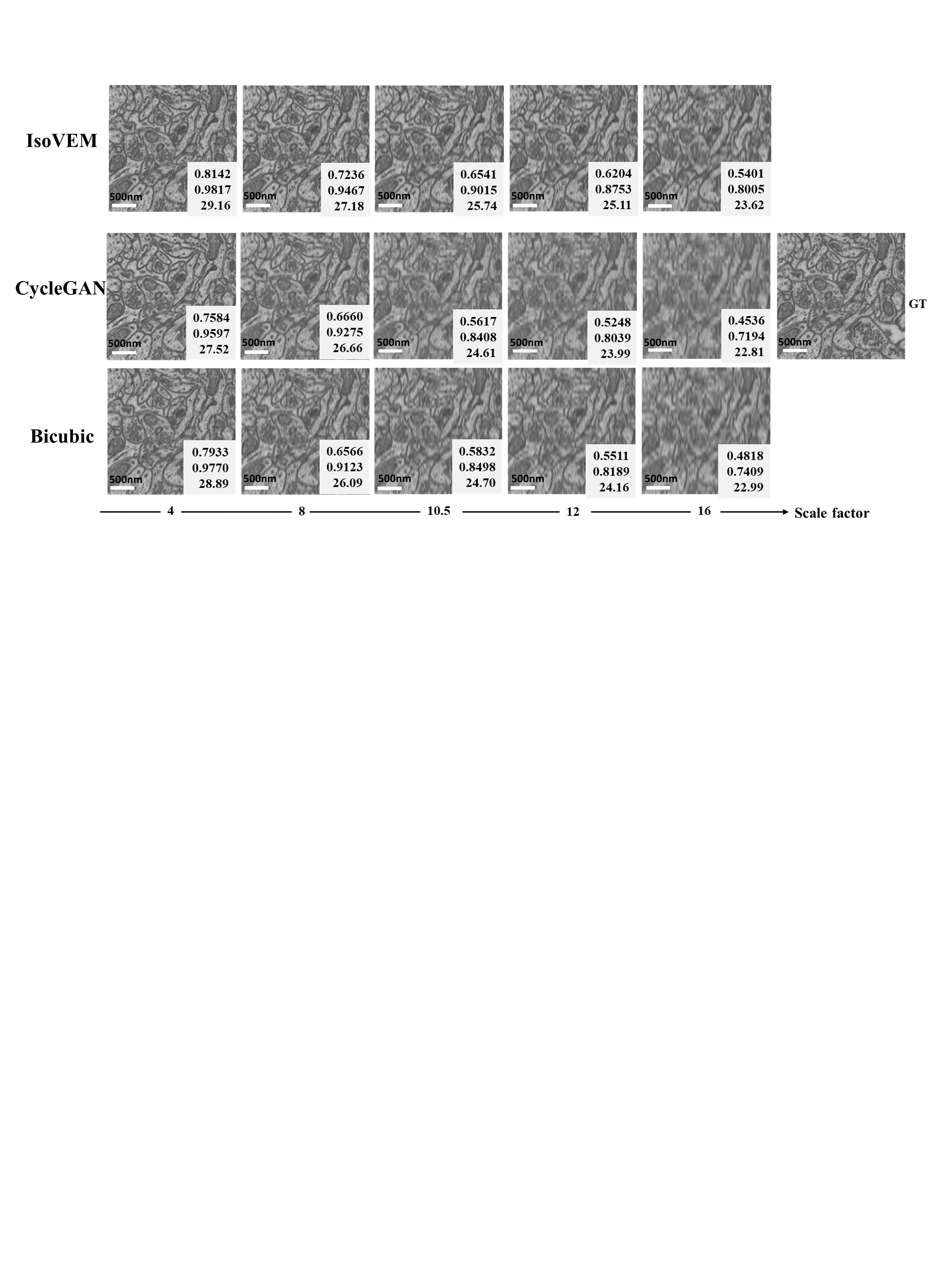

**Extended Data Fig. 7 | Pretrained model inference or interpolation on arbitrary anisotropic simulated EPFL dataset.** For IsoVEM and CycleGAN, we used the model trained on 8x anisotropic simulated EPFL dataset. To realize arbitrary scale reconstruction during inference without changing model weights, IsoVEM takes trilinear interpolation in up-sampling module, while CycleGAN changes the scale factor of up and down sampling module in the network. It can be seen that artifacts appear in CycleGAN due to the mismatch of weight and scale factor, while IsoVEM shows best robustness to scale factor than CycleGAN and bicubic interpolation. The indicators in the lower right corner are SSIM, MS-SSIM and PSNR respectively.
