## Extended Data Figure 8 for "IsoVEM: Isotropic Reconstruction for Volume Electron Microscopy Based on Transformer"

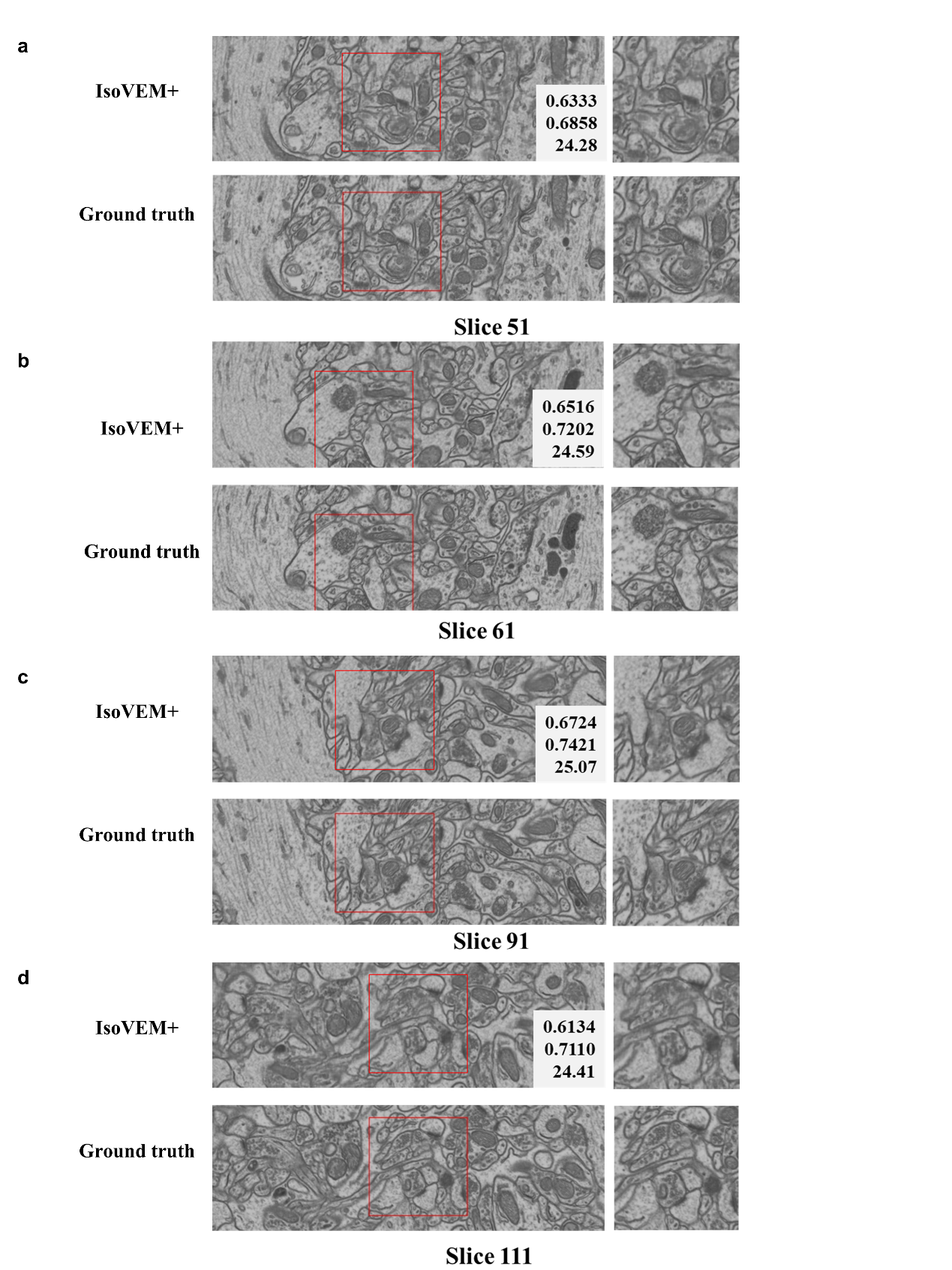

**Extended Data Fig. 8 | Inpainting missing slice using IsoVEM+ on EPFL dataset.** **a,b,c,d,** Comparison of the restored slice between IsoVEM+ and ground truth. Almost all the ultrastructure was restored by IsoVEM+. The indicators in the lower right corner are SSIM, MS-SSIM and PSNR respectively.
