## Supplementary Note 1 for "IsoVEM: Isotropic Reconstruction for Volume Electron Microscopy Based on Transformer"

**Supplementary Note 1. Degradation modeling.**

The general modeling of image degradation is formed as follows. For isotropic reconstruction task of vEM, the key is to modeling down-sampling process along axial direction, while the blurring and noise are not considered in this paper.

(1)

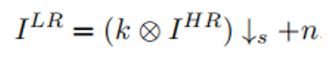

where *I^HR^* is high resolution image, *I^LR^* is low resolution image, *k* is blur kernel, *s* is down-sampling operation, and *n* is noise.

In the sample preparation process for vEM, different techniques are faced with the same problem of information loss in sample thickness direction. This include the information loss caused by serial section thickness in ssSEM and ssTEM technology, and similar information loss caused by the milling thickness in SBF-SEM or FIB-SEM. In addition, the point spread effect during the interaction of the electron beam or ion beam with the camera is also a source of information loss. Regardless of the electron microscope imaging mode, one can generally approximate the imaging process as an integral summation of the sample information along the imaging optical path. If we consider that the electron beam has the same effect on samples at different thicknesses location, the degradation process can be approximated as anisotropic 3D average pooling, which also used by Heinrich^1^. As shown in Fig. 1a, we use anisotropic 3D average pooling to simulate degradation and construct self-supervised training dataset. We further verified that 3d average pooling can be used as a universal representation of the axial degradation of vEM serial slices through transfer learning experiments. For example, the model trained from FIB-SEM has good transferability on ssTEM data, and vice versa. It is worth noting that there are also some vEM isotropic reconstruction works takes frame extraction^2,3^or trainable DNN as degradation process^4^ Our method can be easily extended to this assumption, only changing the degradation operation in the self-supervised data construction.
