## Supplementary Note 2 for "IsoVEM: Isotropic Reconstruction for Volume Electron Microscopy Based on Transformer"

**Supplementary Note 2. Details of Self- and Mutual- attention.**

Define input video clip as , and the super-resolved video clip as . , , are the frame number, height, width of the image respectively, and are the input channel number and output channel number. *s* is the up-sampling scale factor that can be integer or decimal, and it is calculated by anisotropic ratio between the cutting/section thickness and the pixel size for imaging when collect data by vEM.

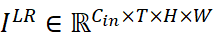

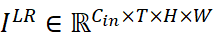

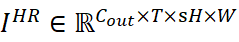

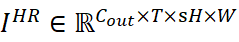

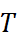

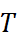

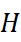

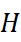

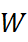

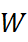

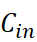

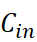

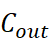

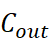

**Self-attention.** Self-attention is a mechanism that makes model decide how important each part of input is, which makes it possible to find dependencies and connections in the data. The computational graph is shown in Supplementary Fig. 1a.

Given a 3D window video clip , where is the number of feature elements and is the channel number. In self-attention, query , key and value are obtained by different linear projection from	itself.

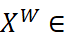

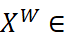

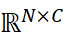

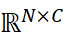

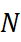

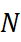

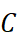

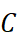

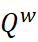

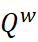

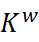

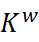

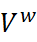

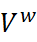

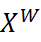

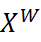

(2)

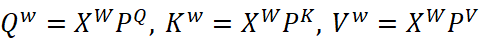

where are projection matrices. is the channel number of projected features.

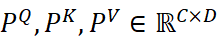

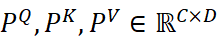

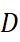

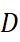

Then the attention map is generated by and , which reflects the importance of different regions.

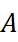

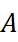

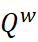

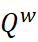

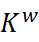

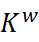

(3)

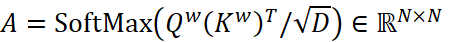

where SoftMax means the row softmax operation.

Finally, the self-attention result is obtained by weighted sum of taking the attention map as the weight.

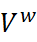

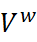

(4)

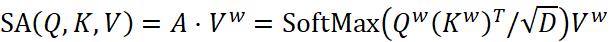

**Mutual attention.** Mutual attention means finding the correlation between elements in two image frames, denote as reference and supporting frames respectively. The computational graph is shown in Supplementary Fig. 1b.

Given a reference frame feature and a supporting frame feature , query is obtained by linear projection of , key and value are obtained by different linear projection of .

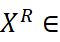

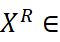

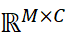

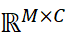

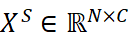

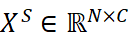

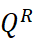

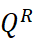

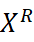

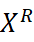

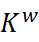

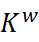

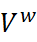

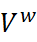

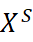

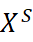

(5)

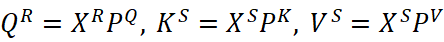

where are projection matrices. is the channel number of projected features. Then, the and are used to generate the attention map . It reflects the correspondence of elements between two frames.

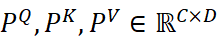

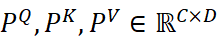

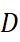

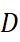

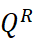

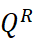

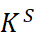

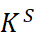

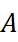

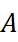

(6)

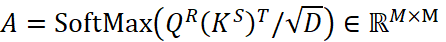

where SoftMax means the row softmax operation.

Finally, the result of mutual attention is generated by weighted sum, which represents new soft-warped feature of the reference frame.

(7)

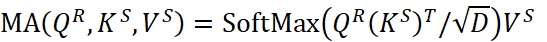

Because comes from support frame, mutual attention result is asymmetric unlike self-attention. Therefore, we also need to exchange the identities of two frames to calculate the other mutual attention between the reference and supporting images.

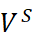

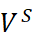

**Shifted window.** Since the complexity of both attentions is quadratic to the number of elements within the attention window, the computation of global attention is impractical. We take shifted window mechanism spatially and temporally to reduce computation time and keep cross-window connections. As shown in Supplementary Fig. 1c, we divide the video clip into non-overlapping local windows and shift it half the length of the partition at next layer of model. Specifically, the window partition in temporal can establish the long-distance dependence of features between frames, one frame can utilize information from up to 2(i − 1) frames at layer i (i ≥ 2). The window partition in spatial make the receptive field size increases as the layer deepens, like Swin Transformer It should be noted that the temporary window size of mutual attention should be 2, but there is no restriction on self-attention.
