## Supplementary figures and images for "IsoVEM: Isotropic Reconstruction for Volume Electron Microscopy Based on Transformer"

### Supplementary Table 1

**Supplementary Table 1. Description of vEM Datasets.**

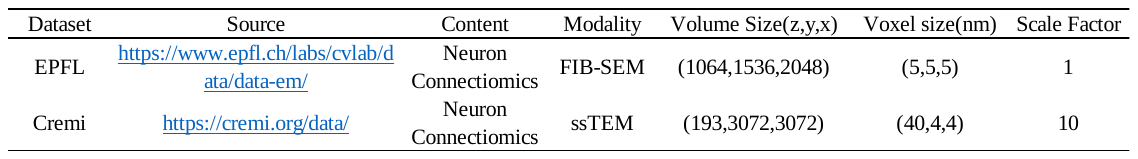
