## Supplementary Table 2 for "IsoVEM: Isotropic Reconstruction for Volume Electron Microscopy Based on Transformer"

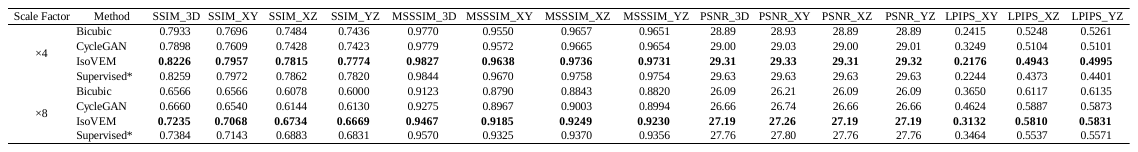

**Supplementary Table 2. Validation metric for isotropic reconstruction results of different methods**

**on EPFL dataset with ground truth.**

**Supplementary Table 4. Transfer Learning of EMformer from Cremi to EPFL dataset with 8-fold anisotropy simulation.**

**Supplementary Table 4. Transfer Learning of EMformer from Cremi to EPFL dataset with 8-fold anisotropy simulation.**
