## Supplementary Table 3 for "IsoVEM: Isotropic Reconstruction for Volume Electron Microscopy Based on Transformer"

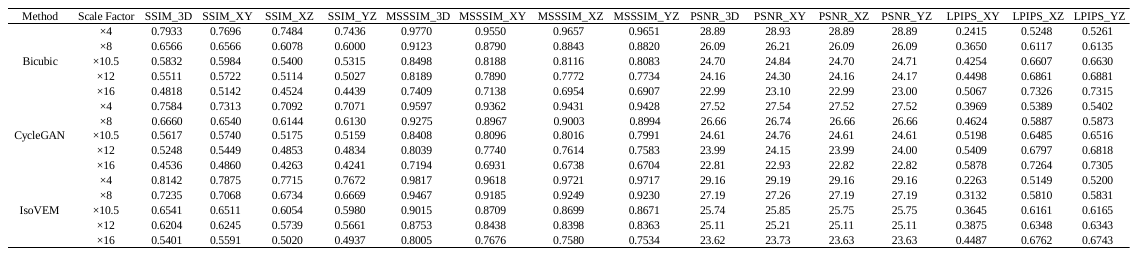

**Supplementary Table 3. Arbitrary scale reconstruction when inference of different method on EPFL dataset.**
