## Supplementary Table 4 for "IsoVEM: Isotropic Reconstruction for Volume Electron Microscopy Based on Transformer"

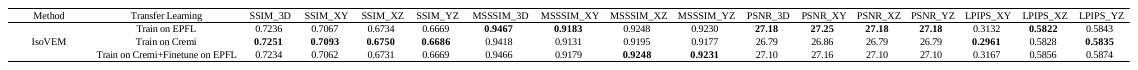

**Supplementary Table 4. Transfer Learning of IsoVEM from Cremi to EPFL dataset with 8X anisotropy simulation.**
