## Supplementary Table 5 for "IsoVEM: Isotropic Reconstruction for Volume Electron Microscopy Based on Transformer"

**Supplementary** **Table 5.** **Validation metric for slice inpainting of IsoVEM+ on EPFL dataset with 8-fold anisotropy simulation.**

**Supplementary Table 4. Transfer Learning of IsoVEM from Cremi to EPFL dataset with 8-fold anisotropy simulation.**

**Supplementary Table 4. Transfer Learning of IsoVEM from Cremi to EPFL dataset with 8-fold anisotropy simulation.**

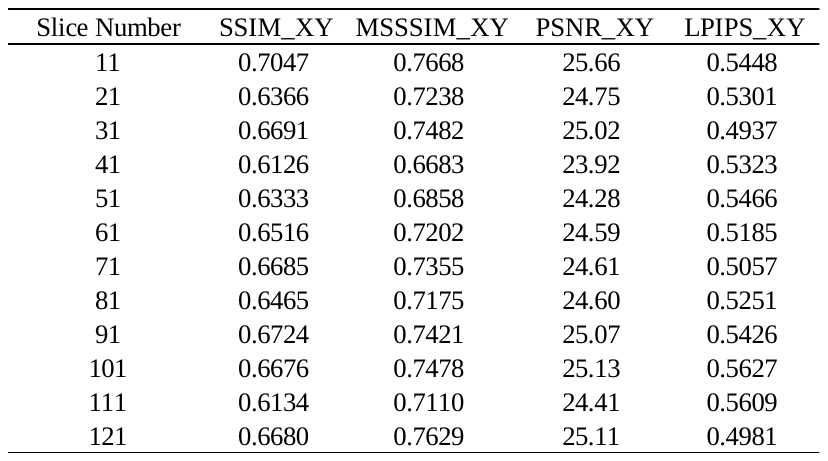
