## Supplementary Table 6 for "IsoVEM: Isotropic Reconstruction for Volume Electron Microscopy Based on Transformer"

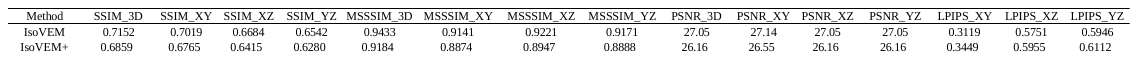

**Supplementary Table 6. Validation metric for isotropic reconstruction of IsoVEM+ with section inpainting on EPFL dataset with 8X anisotropy simulation.**
